# The Arabidopsis PAL3 is a carboxy-tyrosine ammonia lyase

**DOI:** 10.64898/2026.09.15.751672

**Authors:** Caroline Van Beirs, Ans Van der Vaet, Jan Goeman, Fabricio Almeida-Silva, Congwei Xie, Hannes Meinert, Sandrien Desmet, Steven Vandersyppe, Ruben Vanholme, Johan Van der Eycken, Bart Devreese, Thomas Bayer, Uwe T. Bornscheuer, Wout Boerjan, Bartel Vanholme

## Abstract

Phenylalanine ammonia lyase (PAL) catalyses the deamination of L-phenylalanine, which is the first step of the plant-specific phenylpropanoid pathway. Its position at the intersection of primary and specialized metabolism, combined with its important role in plant growth and adaptive stress response, has made it a subject of extensive studies. We identified key amino acids in the catalytic pocket of several PAL enzymes, including PAL3 of Arabidopsis, that are different from the canonical PAL sites, suggesting these enzymes have a different substrate specificity and have been miscategorised for decades. By combining untargeted metabolomics with enzyme assays, we discovered that PAL3 converts 3-carboxy-tyrosine into carboxy-*p*-coumaric acid. Based on this specific function, we propose to denote it as a 3-carboxy-tyrosine ammonia lyase (CAL). In addition to its discovery as a novel enzyme class, paving the way for biotechnological applications, we introduce the carboxy-phenylpropanoids as a new class of specialised metabolites.

## INTRODUCTION

Aromatic amino acid ammonia lyases catalyse the de-amination of aromatic amino acids, yielding the corresponding α,β-unsaturated carboxylic acids and ammonia. Beyond their essential role in maintaining nitrogen homeostasis, the organic acids generated by these enzymes serve as key precursors for a wide range of metabolites involved in growth, defence, and adaptation to environmental stresses^1,2^. Besides their biological role, ammonia lyases are of significant interest in green chemistry due to their ability to catalyse amination reactions. This activity facilitates the synthesis of aromatic compounds and stereospecific natural and non-natural amino acids, which serve as valuable precursors for pharmaceuticals and other industrially relevant chemicals^1,3,4^.

Based on their substrate specificity, aromatic amino acid ammonia lyases are classified into three groups: phenylalanine ammonia lyases (PALs; EC4.3.1.24) that convert L-phenylalanine to cinnamic acid, tyrosine ammonia lyases (TALs; EC4.3.1.23) that deaminate L-tyrosine to *p*-coumaric acid, and histidine ammonia lyases (HALs; EC4.3.1.3) that convert L-histidine to urocanic acid (**Fig. 1A**). While HALs are present in animals and microorganisms but absent in plants, PALs and TALs are found in microorganisms and plants, but not in animals^1^. Interestingly, TAL enzymes of plants also have PAL activity and are consequently named phenylalanine/tyrosine ammonia lyases^5^ (PTALs). So far, these PTALs have only been found in grasses, in contrast to PALs which are universally present across all plants^5^.

**Figure 1:**
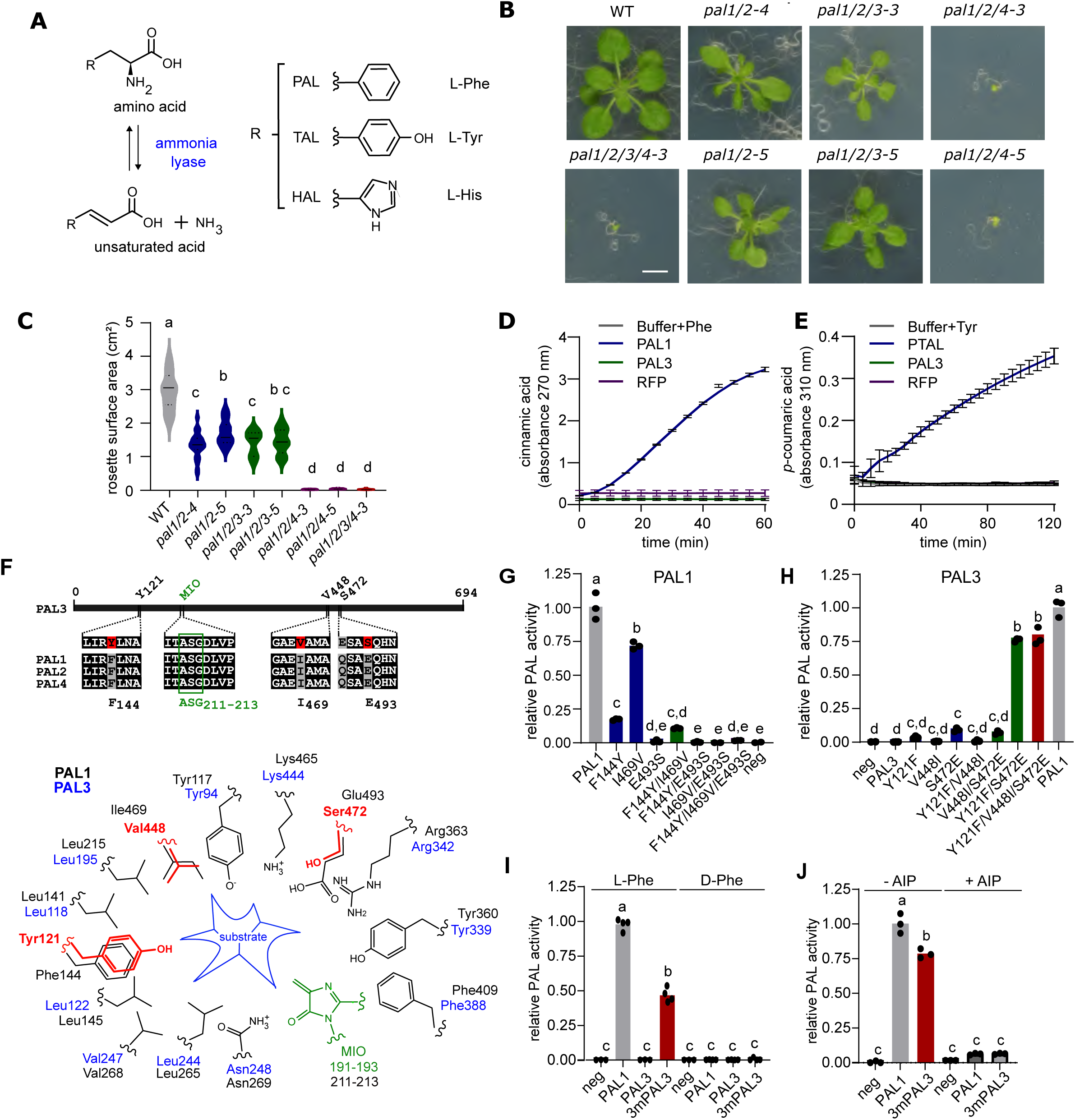
Arabidopsis PAL3 is an ammonia lyase with a noncanonical active site. (**A**) Enzymatic reaction of the three known 4-methylideneimidazol-5-one (MIO)-prosthetic group-dependent aromatic ammonia lyases. (**B**) Rosettes from higher order Arabidopsis *pal* mutants 21 DAS. Plants were grown on ½ MS medium under long day growth conditions (16 h light / 8 h dark) at 21 °C. Scale bar indicates 1 cm. (**C**) Violin plot of rosette surface areas from higher order *pal* mutants shown in (B). Statistical significances were determined using an Anova test followed by a Tukey post hoc test (*p* ≤ 0.05). Equal letters indicate no statistical significance. (**D**) PAL enzyme assay using proteins heterologously produced in *E. coli*. L-phenylalanine was used as substrate. Absorbance at 270 nm was used as a proxy of cinnamic acid production. PAL1 was included as a positive control, RFP was included as a negative control. n =3. (**E**) TAL enzyme assay using proteins heterologously produced in *E. coli*. L-tyrosine was used as substrate. Absorbance at 310 nm was used as a proxy of *p*-coumaric acid production. PTAL from *Brachypodium distachyon* was included as a positive control, RFP was included as a negative control. n = 6. (**F**) Linear representation of Arabidopsis PAL sequences. Amino acid residues differing between PAL1/2/4 and PAL3 are highlighted in red. The ASG tripeptide forming the MIO-group is indicated in green. The catalytic pockets of Arabidopsis PAL1 and PAL3 are illustrated below the alignment. Distinct amino acid residues between both sequences are indicated in red. The MIO-group is indicated in green. (**G**-**J**) PAL enzyme assays as in (D) with L-phenylalanine as substrate. Absorbance at 270 nm was used as a proxy of cinnamic acid production, and absorbance values were normalized to PAL1. (**G**) focuses on different mutated variants of PAL1 whereas (**H**) focuses on different mutated variants of PAL3. (**I**) and (**J**) focus on PAL1 and triple mutated PAL3_Y121F/V448I/S472E_ (3mPAL3). In (I) the effect of the ammonia lyase inhibitor AIP was investigated whereas in (J) the stereospecificity of the enzyme was tested by using both L-phenylalanine or D-phenylalanine as substrate. AIP: 2-amino-2-inden-phosphonic acid, DAS: days after stratification, GC-MS: gas chromatography-mass spectrometry, HAL: histidine ammonia lyase, MIO: 4-methylideneimidazol-5-one, PAL: phenylalanine ammonia lyase, Phe: L-phenylalanine, RFP: red fluorescent protein, TAL: tyrosine ammonia lyase, WT: wildtype

In plants, PALs and PTALs catalyse the initial step of the phenylpropanoid pathway, producing the precursor for a wide range of phenolic metabolites essential to diverse physiological processes, thereby playing a critical role in plant growth and development^2^. Among the phenylpropanoid pathway-derived compounds are the building blocks for lignin, the earth’s most abundant plant polymer after cellulose. By providing mechanical strength to the plant cell wall, lignin enables plants to grow upright and withstand various environmental stresses, including wind and water pressure^6^. Additional phenylpropanoid pathway-derived compounds that depend on PAL and PTAL for their synthesis are coumarins, flavonoids, lignans, and stilbenes^7^. Many of these compounds play a vital role in protecting the plant against biotic and abiotic stresses.

*Arabidopsis thaliana* has four homologous *PAL* genes (*AtPAL1-4*; AT2G37040, AT3G53260, AT5G04230, and AT3G10340; hereafter referred to as *PAL1-4*), which exhibit both redundant and specialized functions in development and metabolism^8–10^. For example, *PAL1* and *PAL2* are involved in lignification and defence against (a)biotic stresses^9,10^, whereas *PAL4* has been linked to suberization^11,12^. *PAL3* stands out as the least studied member of the family, as no enzymatic evidence currently supports its PAL activity, and the gene is scarcely represented in expression databases^13^. Recently, the aberrant behaviour of PAL3 compared to other PALs in Arabidopsis was highlighted again, and it was suggested that PAL3 may have a distinct, yet unidentified, biological function^14^. We performed a detailed analysis of multiple sequence alignments which revealed critical amino acid differences in the catalytic pocket of PAL3, and suggesting a deviating substrate specificity from the traditional PALs. By combining untargeted metabolomics of Arabidopsis *pal3* mutants with in-depth enzyme analyses, we discovered that PAL3 does not use L-phenylalanine as substrate, but converts 3-carboxy-tyrosine into carboxy-*p*-coumaric acid.

## RESULTS

### Arabidopsis PAL3 has no PAL activity

To explore the enigmatic role of *PAL3* and assess potential functional redundancy with the three better characterized *PAL* genes in Arabidopsis, CRISPR mutants were generated for each of the four genes. Gene-specific gRNAs were designed against the exon sequence upstream or in the region coding for the catalytic pocket. For each of the four *PAL* genes, two independent CRISPR lines with a frameshift mutation were selected (**Supplementary Fig. 1**). The *pal1/2* double mutants were generated via a multiplex CRISPR strategy, the other higher order mutants (i.e. *pal1/2/3*, *pal1/2/4* and *pal1/2/3/4*) were obtained by crossing mutant lines followed by selfing. Sequence-verified homozygous mutant plants were grown on horizontal plates containing 1/2 MS medium, and the rosettes were analysed 21 days after stratification (DAS). Whereas none of the single mutants showed a visual reduction in rosette size compared with wild-type (WT) (**Supplementary Fig. 2**), *pal1/2-4* and *pal1/2-5* double mutants showed a 54% and 42% reduction in rosette surface areas (**Fig. 1B, C**). Introducing the *pal3* mutation in the *pal1/2* mutant background did not further affect the rosette surface area. However, introducing a *pal4* mutation in the *pal1/2* mutant background resulted in rosette sizes of 0.65% and 1.1% as compared with WT for *pal1/2/4-3* and *pal1/2/4-5*, respectively. A similar dramatic decrease in rosette size was observed in the *pal1/2/3/4-3* quadruple mutant (0.61% compared with WT). In summary, this experiment demonstrated no obvious redundancy between *PAL3* and *PAL1, 2* or *4* at the phenotypic level.

To investigate whether PAL3 has PAL activity, PAL3 was heterologously produced in *Escherichia coli* as a His-MBP-tagged fusion protein (**Supplementary Fig. 3**). After protein purification, an enzyme assay was performed using L-phenylalanine as substrate, and the enzymatic activity was quantified by measuring the absorbance of the PAL product cinnamic acid spectrophotometrically at 270 nm. PAL1, which was included as a positive control, showed clear production of cinnamic acid over time, indicating PAL activity (**Fig. 1D**). No formation of cinnamic acid was observed when no protein was added to the assay, or when PAL1 was replaced by the non-enzymatic red fluorescent reporter protein (RFP). Similarly, no conversion of L-phenylalanine to cinnamic acid was detected using PAL3, indicating the enzyme has no PAL activity. In parallel, PAL3 was tested for activity towards L-tyrosine, indicative of TAL activity. The enzymatic activity was quantified by measuring absorbance of the TAL product *p*-coumaric acid spectrophotometrically at 310 nm. A heterologously produced PTAL enzyme of *Brachypodium dystachion,* included as a positive control^5^, showed clear conversion to *p*-coumaric acid over time (**Fig 1E**). No production of *p*-coumaric acid was observed for PAL3, indicating PAL3 does not have TAL activity.

### PAL3 is an ammonia lyase with a noncanonical active site

Despite the absence of activity against L-phenylalanine and L-tyrosine, reflecting the ammonia lyase activities known to plants, PAL3 possesses the alanine-serine-glycine (ASG191-193) tripeptide that forms the 4-methylideneimidazol-5-one (MIO)-prosthetic group characterizing the catalytic pocket of ammonia lyases^1,15^ (**Fig. 1F; Supplementary Fig. 4**). This indicates PAL3 is likely an ammonia lyase, although acting on a substrate different from L-phenylalanine or L-tyrosine. The substrate specificity of ammonia lyases was reported to be determined by a single amino acid in the catalytic pocket (F144 for Arabidopsis PAL1), and is known as the substrate specificity site in literature^16^. This crucial amino acid is a histidine in (P)TALs, a serine in HALs, and a phenylalanine in PALs. Intriguingly, PAL3 has a tyrosine at this position (Y121), supporting the absence of PAL and TAL activity, but also suggesting that PAL3 does not have HAL activity. Comparison of the catalytic pocket of PAL3 with that of PAL1 revealed two additional amino acid differences: isoleucine to valine at position 448 (V448), and glutamic acid to serine at position 472 (S472; **Fig. 1F**).

Building on this observation, variants of PAL1 were generated by exchanging these three amino acids with the corresponding residues from PAL3, including single substitutions (PAL1_F144Y,_ PAL1_I469V_ and PAL1_E493S_), as well as double (PAL1_F144Y/I469V,_ PAL1_F144Y/E493S_ and PAL1_I469V/E493S_) and triple (PAL1_F144Y/I469V/E493S_) combinations. These enzymes were produced heterologously in *E. coli* and assayed as described before (**Supplementary Fig. 3**). PAL1, included as a positive control, showed strong PAL activity, validating the reliability of the experimental setup (**Fig. 1G**). A significant reduction in PAL activity was observed for PAL1_F144Y,_ confirming the key role of this site in determining substrate specificity and catalytic turnover. However, the observation that the loss of activity is only partial indicates that additional amino acid residues contribute to the configuration and function of the catalytic pocket. Mutating the second site (I469V) also impacted PAL activity, but to a lesser extent. Combining both mutations did not have an additional negative effect on PAL activity, as the activity of the single PAL1_F144Y_ mutant was similar to that of the PAL1_F144Y/I469V_ double mutant. Interestingly, of the three differing amino acids under study here, E472S had the biggest impact on PAL activity, as the PAL1_E493S_ single mutant completely lost its PAL activity. In line with this result, neither the double (PAL1_F144Y/E493S,_ PAL1_I469V/E493S_) nor the triple (PAL1_F144Y/I469V/E493S_) mutants containing this mutation showed any residual PAL activity.

Conversely, variants of PAL3 were generated by introducing the corresponding PAL1 amino acids, using single, double and triple substitutions with the aim of identifying key amino acids that are capable of introducing PAL activity in PAL3 (**Fig. 1H; Supplementary Fig. 3**). Consistent with previous results, PAL3 WT did not display activity towards L-phenylalanine. No significant enhancement in PAL activity was detected following individual site-directed substitution of Y121F or V448I. In contrast, a small but significant increase in enzyme activity was observed upon introduction of the S472E mutation. When this mutation was combined with Y121F, a remarkable increase in PAL activity was observed, resulting in activity levels of ∼75% relative to PAL1. Introducing V448I into this double mutant did not further increase PAL activity. Analogous to PAL1, this PAL3 variant (i.e. 3mPAL3) was stereospecific, as only L-phenylalanine and not D-phenylalanine could be converted to cinnamic acid (**Fig 1I**). Additionally, this triple mutated enzyme variant was susceptible to inhibition using the PAL inhibitor 2-amino-2-inden-2-phosphonic acid^17,18^ (AIP; **Fig 1J**).

As the primary substrates of known aromatic amino acid ammonia lyases are amino acids (i.e. L-phenylalanine, L-tyrosine or L-histidine), PAL3 activity was assayed on a mixture of 20 proteogenic amino acids, where substrate conversion should result in a decrease in abundance of the corresponding amino acid. All samples were incubated for 30 minutes and consequently analysed using gas chromatography-mass spectrometry (GC-MS). Of the 20 amino acids used in the assay, arginine and alanine could not be detected with the applied method. Incubating the 20 amino acids with PAL1, which was included as a positive control, significantly decreased L-phenylalanine content without affecting the levels of the other amino acids, indicative of PAL activity (Supplementary Fig. **Fig. 5**). In contrast, no significant decrease in amino acid abundance was observed for PAL3, indicating that PAL3 does not take any of these amino acids as substrate.

Together, these data demonstrate that besides the known substrate specificity site (Y121) of ammonia lyases, two other positions in the catalytic pocket (V448 and S472) contribute to PAL activity and/or specificity. Additionally, we showed that PAL activity can be introduced in PAL3, indicating that - despite the fact that WT PAL3 does not possess PAL activity - intrinsic ammonia lyase properties are preserved in this enzyme.

### *pal3* mutants have a metabolic phenotype

To identify the substrate(s) and product(s) of PAL3, we employed a metabolomics approach using the generated *pal3-3* and *pal3-4* CRISPR mutants. The underlying principle is that, in absence of complete genetic redundancy, substrate(s) and product(s) of PAL3 are expected to accumulate and decrease, respectively, in *pal3* mutants. To determine the appropriate tissue and developmental stage for metabolomic analysis, we first examined the expression pattern of *PAL3*. A general low expression of *PAL3* is reported by the eFP browser 2.0 (**Supplementary Fig. 6A**). To obtain a more detailed image of *PAL3* expression in the plant, transcriptional β-glucuronidase (*pPAL3::GUS*) reporter lines were generated using a 2.2-kb genomic region upstream of the *PAL3* translation initiation site. The reporter lines were grown *in vitro* on vertical plates containing 1/2 MS medium. At 7 DAS, *PAL3* promoter activity was found mainly in the roots, and not found in the cotyledons (**Supplementary Fig. 6B-D**). At 14 DAS, *PAL3* promoter activity remained present in the roots, and to a lesser extent in the vasculature of the leaves (**Supplementary Fig. 6E-G**). Based on these results, roots harvested at 14 DAS were selected for subsequent analysis.

Root methanol extracts were analysed via reverse-phase ultra-high-performance liquid chromatography–mass spectrometry (UHPLC-MS). In total, 18,468 *m*/*z* features were detected and integrated in each of the samples. Statistical analysis and applying the requirement of a fold-change ratio of at least two between WT and both *pal3* mutants finally resulted in a list of differential *m*/*z* signals. Eleven were reduced in intensity in both *pal3-3* and *pal3-4*, as compared to WT (**Table 1**), making them potential downstream PAL3 products. Based on MS/MS spectral fragmentation, the structure of ten of these compounds could be elucidated as carboxy-phenylpropanoids. The use of a chemically synthesised reference compound indicated that all ten compounds belonged to a newly discovered metabolic class of carboxy-phenylpropanoids (**Table 1**, **Fig. 2, Supplemental Fig. 7**). Compound **1** matched with the carboxy-*p*-coumaric acid reference compound, whereas compounds **2** to **6** appeared to be derivates thereof: **2** and **3** were characterized as carboxy-*p*-coumaric acid linked to a hexose moiety, **4** and **5** appeared to be carboxy-*p*-coumaric acid linked to a lysine moiety, and **6** was characterized as carboxy-*p*-coumaric acid linked to putrescine. Compound **7** and **8** were characterized as hexose derivates of carboxy-caffeic acid, the latter presumably being a hydroxylation product of carboxy-*p*-coumaric acid. Finally, compound **9** and **10** could be structurally elucidated as carboxy-ferulic acid, i.e., the methylated product of carboxy-caffeic acid. In addition, two *m*/*z* features appeared to be increased by at least two-fold in both *pal3-3* and *pal3-4* samples (**12** and **13**, **Table 1**), suggesting that these features may represent putative substrates of PAL3 or derivates thereof. Unfortunately, we could not elucidate the structures of the corresponding compounds.

**Figure 2:**
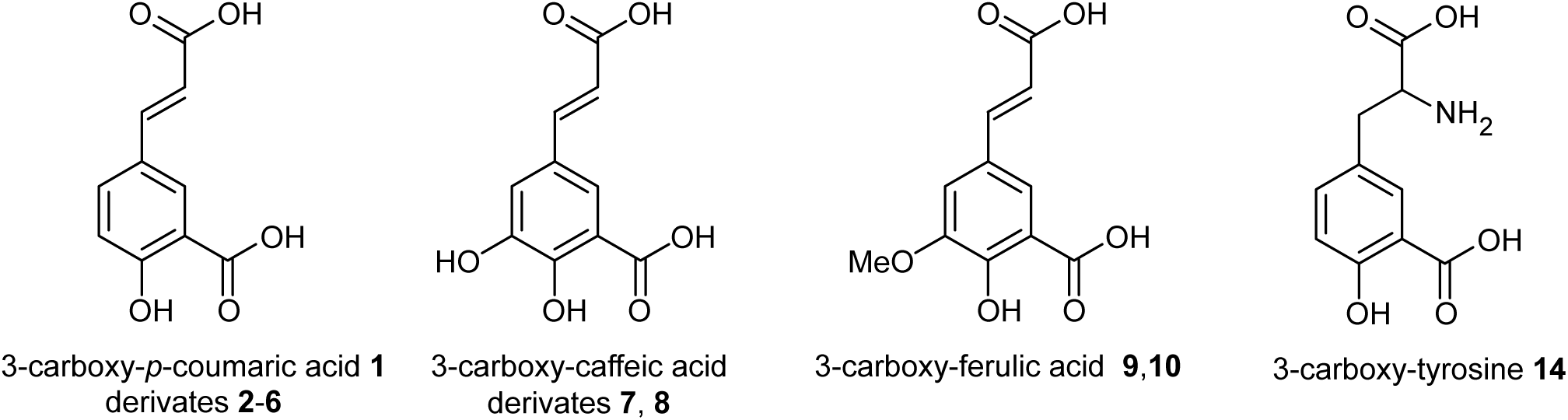
Differentially accumulating carboxy-phenylpropanoids in roots of 14 DAS *pal3* mutants. Structure of the compounds presented in Table 1. For structural elucidation of the different molecules see Supplemental Figure S4.

**Table 1:** Differentially accumulating 3-carboxy-phenylpropanoids in roots of 14 DAS *pal3* mutants. Comparative metabolomics data from *pal3* mutants and WT. Compounds **1**-**13** are retrieved via untargeted metabolomics. Compounds **14**-**15**, indicated with a *, were retrieved via a targeted approach. Average values represent normalized peak intensity. RT: retention time. The structures of the characterized metabolites are shown in Figure 2.

| No. | Name | RT (min) | $m/z_{\text{exp}}$ | WT<br>average ± s.d. | *pal3-3*<br>average ± s.d. | fold change<br>*pal3-3*/WT | *pal3-4*<br>average ± s.d. | fold change<br>*pal3-4*/WT |
| --- | --- | --- | --- | --- | --- | --- | --- | --- |
| *reduced abundance in pal3 mutants* |  |  |  |  |  |  |  |  |
| 1 | 3-carboxy-*p*-coumaric acid | 13.53 | 207.0287 | 68200 ± 3100 | 9730 ± 1120 | 0.14 | 12100 ± 2680 | 0.18 |
| 2 | 3-carboxy-*p*-coumaric acid + hexose 1 | 10.14 | 369.0822 | 50100 ± 12000 | 5030 ± 2470 | 0.10 | 3770 ± 2120 | 0.08 |
| 3 | 3-carboxy-*p*-coumaric acid + hexose 2 | 10.54 | 369.0812 | 54500 ± 15700 | 16400 ± 4960 | 0.30 | 15200 ± 4130 | 0.28 |
| 4 | 3-carboxy-*p*-coumaric acid + lysine 1 | 8.21 | 335.1242 | 11600 ± 5000 | 5 ± 12 | 0.00 | 0 ± 0 | 0.00 |
| 5 | 3-carboxy-*p*-coumaric acid + lysine 2 | 8.97 | 335.1281 | 17300 ± 3196 | 8000 ± 1660 | 0.46 | 5710 ± 3290 | 0.33 |
| 6 | 3-carboxy-*p*-coumaric acid + putrescine | 8.17 | 277.1206 | 109000 ± 17000 | 40200 ± 6470 | 0.37 | 36600 ± 10400 | 0.34 |
| 7 | 3-carboxy-caffeic acid + hexose 1 | 5.85 | 385.0764 | 182000 ± 24400 | 23900 ± 6680 | 0.13 | 19000 ± 11400 | 0.10 |
| 8 | 3-carboxy-caffeic acid + hexose 2 | 7.33 | 385.0766 | 124000 ± 16100 | 9790 ± 2600 | 0.08 | 8030 ± 4340 | 0.06 |
| 9 | 3-carboxy-ferulic acid 1 | 12.07 | 237.0386 | 84000 ± 22700 | 18900 ± 10800 | 0.23 | 16100 ± 6590 | 0.19 |
| 10 | 3-carboxy-ferulic acid 2 | 12.73 | 237.0387 | 42000 ± 2170 | 7010 ± 442 | 0.17 | 8950 ± 2150 | 0.21 |
| 11 | unkown 1 | 21.8 | 328.0602 | 25600 ± 4930 | 4130 ± 1560 | 0.16 | 12500 ± 2160 | 0.49 |
| *increased abundance in pal3 mutants* |  |  |  |  |  |  |  |  |
| 12 | unkown 2 | 2.21 | 812.9835 | 20400 ± 7350 | 85500 ± 19900 | 4.19 | 638900 ± 19500 | 31.32 |
| 13 | unkown 3 | 2.58 | 662.102 | 10000 ± 3780 | 40600 ± 3310 | 4.06 | 26600 ± 5940 | 2.66 |
| 14 | 3-carboxy-tyrosine * | 4.38 | 224.0552 | 3268000 ± 172000 | 3980000 ± 138000 | 1.22 | 4450000 ± 90800 | 1.36 |
| *not differential* |  |  |  |  |  |  |  |  |
| 15 | 3-carboxy-phenylalanine * | 4.42 | 208.0601 | 2050000 ± 151000 | 2000000 ± 85400 | 0.98 | 2090000 ± 83700 | 1.02 |

In an attempt to identify potential substrates of PAL3, we switched to a targeted approach. Given the identification of the carboxy-phenylpropanoids as potential downstream products of PAL3, combined with the ammonia lyase activity of PAL3, we considered 3-carboxy-phenylalanine or 3-carboxy-tyrosine as potential substrate(s) of PAL3. Whereas the targeted analysis showed no change in the levels of 3-carboxy-phenylalanine (**15**), 3-carboxy-tyrosine (**14**) was increased in both *pal3* lines (p-value 0.0016 and 0.0001 in *pal3-3* and *pal3-4*, respectively; **Table 1**). Given the relative increase in 3-carboxy-tyrosine was moderate (22% and 36%, respectively), it had not triggered the requirement of ‘at least two-fold increase’ used in the initial untargeted analysis.

### PAL3 is a carboxy-tyrosine ammonia lyase

Based on the metabolic profiles of *pal3* mutants, we speculated that PAL3 could possess carboxy-tyrosine ammonia lyase (CAL) activity, converting 3-carboxy-tyrosine into carboxy-*p*-coumaric acid and ammonia (**Fig. 3A**). To validate this hypothesis, enzyme assays using heterologously produced PAL3 and PAL1 were performed with chemically synthesized 3-carboxy-tyrosine as the substrate, and samples were analysed using high performance liquid chromatography mass spectrometry (HPLC-MS). No 3-carboxy-*p*-coumaric acid was found in the negative control, indicating that 3-carboxy-tyrosine does not spontaneously convert to carboxy-*p*-coumaric acid (**Fig. 3B, C**). Likewise, no 3-carboxy-*p*-coumaric acid was detected when PAL1 was supplied with 3-carboxy-tyrosine, indicating PAL1 does not possess CAL activity. In contrast, PAL3 was able to convert 3-carboxy-tyrosine into carboxy-*p*-coumaric acid and the conversion could be suppressed by the addition of the ammonia lyase inhibitor AIP ^17,18^ **Fig. 3D**). Pronounced substrate inhibition was observed at concentrations above 500 µM 3-carboxy-tyrosine (**Supplementary Fig. 8**).

**Figure 3:**
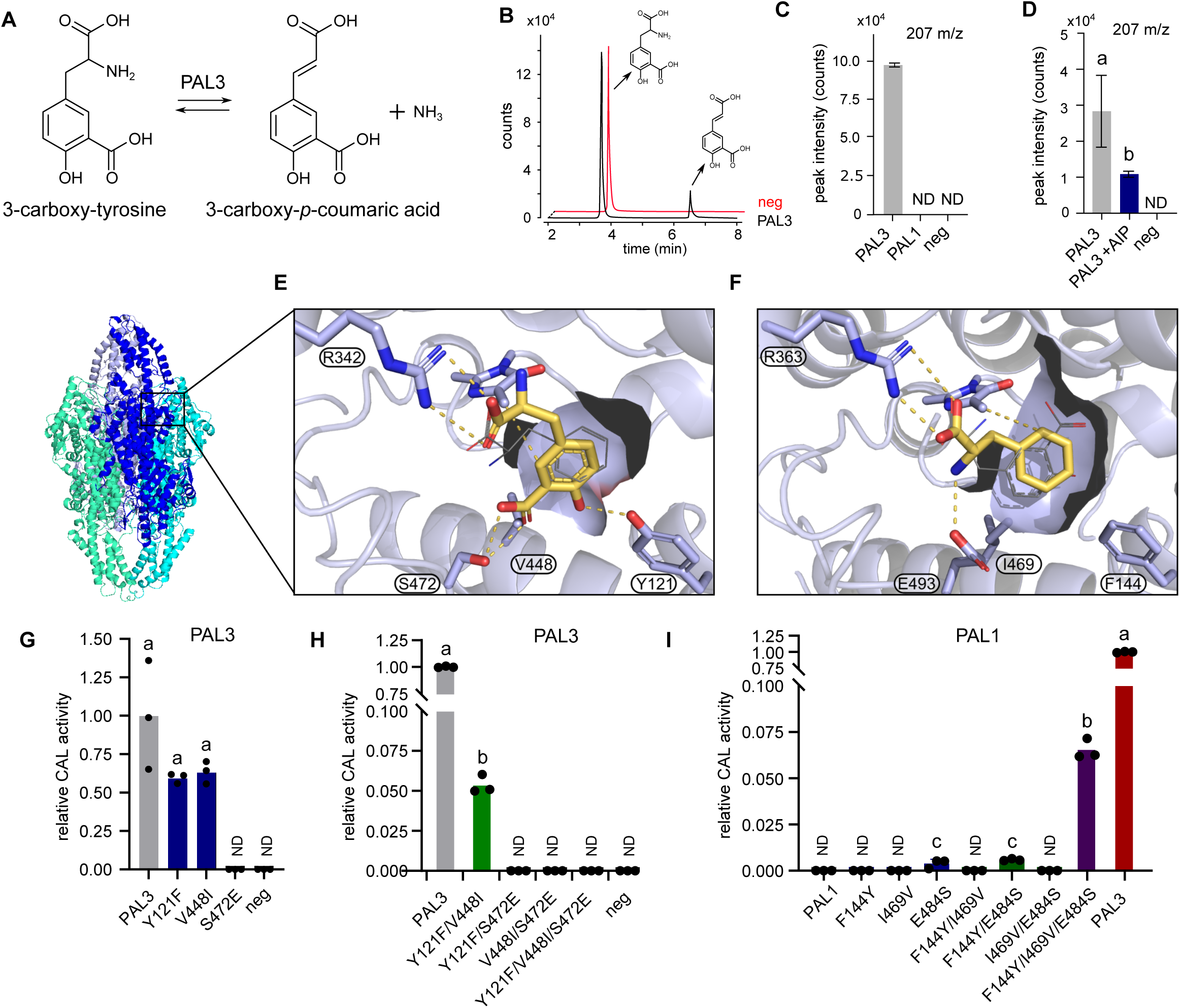
Arabidopsis PAL3 is a 3-carboxy-tyrosine ammonia lyase. (**A**) Enzymatic reaction catalysed by PAL3 whereby 3-carboxy-tyrosine is converted into 3-carboxy-*p*-coumaric acid. (**B-D**) Enzyme assays using proteins heterologously produced in *E. coli* and 3-carboxy-tyrosine as substrate. No protein was added in the negative control. Product formation was analysed using HPLC-MS. Statistical significances were determined using an Anova test followed by a Tukey post hoc test (*p* ≤ 0.05). Equal letters indicate no statistical significance. (**B**) shows a representative chromatogram of separated samples with 3-carboxy-tyrosine and carboxy-*p*-coumaric acid, negative control indicated in red and sample with PAL3 indicated in black. (**C**) Compares the activity of PAL1 and PAL3, while (**D**) investigate the effect of the ammonia lyase inhibitor AIP on the conversion efficiency of PAL3 on 3-carboxy-tyrosine. (**E**) Catalytic pocket of PAL3 docked with 3-carboxy-tyrosine (yellow model) or L-phenylalanine (grey stick model). (**F**) Catalytic pocket of PAL1 docked with L-phenylalanine (yellow model) or 3-carboxy-tyrosine (grey stick model). (**G-I**) PAL3 enzyme assays as in (C), with product formation detected by MS/MS was normalized to the corresponding substrate amount detected by MS/MS and subsequently normalized to the PAL3 positive control. In (**G**) the activity of mutated variants of PAL3 were tested using 0.4 mM 3-carboxy-tyrosine as substrate, whereas in (**H**) 1 mM 3-carboxy-tyrosine was used. In (**I**) mutated variants of PAL1 were tested with 0.4 mM 3-carboxy-tyrosine as substrate. AIP: 2-amino-2-inden-2-phosphonic acid, CAL: carboxy-tyrosine ammonia lyase, HPLC-MS: high performance liquid chromatography mass spectrometry, PAL: phenylalanine ammonia lyase, ND: not detected.

Having confirmed PAL3’s preference for 3-carboxy-tyrosine, we aimed to explore how the previously identified amino acid residues interact with and discriminate between 3-carboxy-tyrosine and L-phenylalanine. Protein models were created using the AlphaFold3 server, and Autodock Vina was used to introduce 3-carboxy-tyrosine or phenylalanine into the catalytic pocket of PAL3 and PAL1. The catalytic pocket of PAL1 has a hydrophobic section to which residues F144 and I469 contribute, to accommodate the aromatic ring of L-phenylalanine (**Fig. 3E**). This hydrophobic pocket of PAL1 likely precludes binding of the more hydrophilic ring of 3-carboxy-tyrosine. On the other hand, E493 might coordinate the α-amino group, supporting a Friedel-Crafts reaction mechanism. Conversely, docking of 3-carboxy-tyrosine into the catalytic pocket of PAL3 reveals that S472 might coordinate the negatively charged 3-carboxyl group on the aromatic ring of the substrate, while its hydroxyl group is coordinated by Y121 (**Fig. 3F**). In contrast, binding of L-phenylalanine in PAL3 is hindered by the presence of Y121, which disrupts the hydrophobic pocket needed for proper accommodation of L-phenylalanine.

To elucidate the relative contribution of the three different amino acid residues as outlined by the *in silico* docking on the catalytic activity of PAL3, the previously generated single, double and triple mutants were tested for CAL activity. Mutating Y121F or V448I in PAL3 had a mild effect on the activity (**Fig. 3G**), and combining both mutations (Y121F/V448I) similarly impacted CAL activity (**Fig. 3H**). However, mutating S472E alone, or in combination with the other two amino acid sites, completely abolished CAL activity. By analogy with the amino acid substitutions used to convert PAL3 into a PAL enzyme, we mutated the equivalent positions in PAL1 (**Fig. 3I**) with the respective amino acids from PAL3 to assess whether CAL activity could be introduced into PAL1. No CAL activity was observed for F144Y, I469V or their combination. A mild CAL activity was observed for E493S and the double mutant F144Y/E493S. The combination of all three mutations resulted in the highest CAL activity of all PAL1 variants.

Together, these results confirm the discovery of the native activity of PAL3, converting 3-carboxy-tyrosine into 3-carboxy-*p*-coumaric acid. Among the three characteristic amino acids in the catalytic pocket of PAL3, S472 was identified as the key residue determining substrate specificity, and to a lesser extent Y121.

### PAL3 in the tree of life

The discovery of a novel type of ammonia lyase in *A. thaliana* raised the question of whether PAL3 proteins are unique to this species or conserved across other plant taxa. To explore this, we used delineated gene family assignments from the PLAZA database to identify PAL homologs across plant genomes. Among the 131 plant species screened, comprising monocots, dicots, bryophytes and green algae, a total of 1179 putative ammonia lyase proteins (i.e. PALs, PTALs and PAL3s) were identified. Of these ammonia lyases, only two sequences retain all three PAL3-specific sites, namely those from *A. thaliana* and *A. lyrata* (**Fig. 4A, B; Supplementary Fig. 9**). To assess whether sequences with PAL3-specific sites are restricted to the *Arabidopsis* genus, we identified genes encoding ammonia lyases on UniProt genomes (some of which unavailable on PLAZA). Our analyses revealed three more species containing the PAL3 triad, namely proteins from *A. thaliana x arenosa* (Uniprot code A0A8T2CNU9), *A. suecica* (A0A8T1Z666) and *Arabis nemorensis* (A0A565CHX8) (**Supplementary Fig. 10A**). These proteins are highly conserved, with the least similar pair being *A. thaliana* PAL3 and *Arabis nemorensis* PAL (93.7% similarity and 87.7% identity; **Supplementary Fig 10B.**) Considering the topology of the Brassicaceae phylogeny^19^, parsimony indicates that the PAL3 triad evolved two times independently, once in *A. nemorensis*, and once in the *Arabidopsis* genus (**Supplementary Fig. 11**). The *A. nemorensis* PAL protein was produced heterologously in *E. coli* as described before, and *in vitro* enzyme assays revealed that this enzyme did not have PAL or TAL activity, but instead was able to convert carboxy-tyrosine (**Fig. 4C-E**).

**Figure 4:**
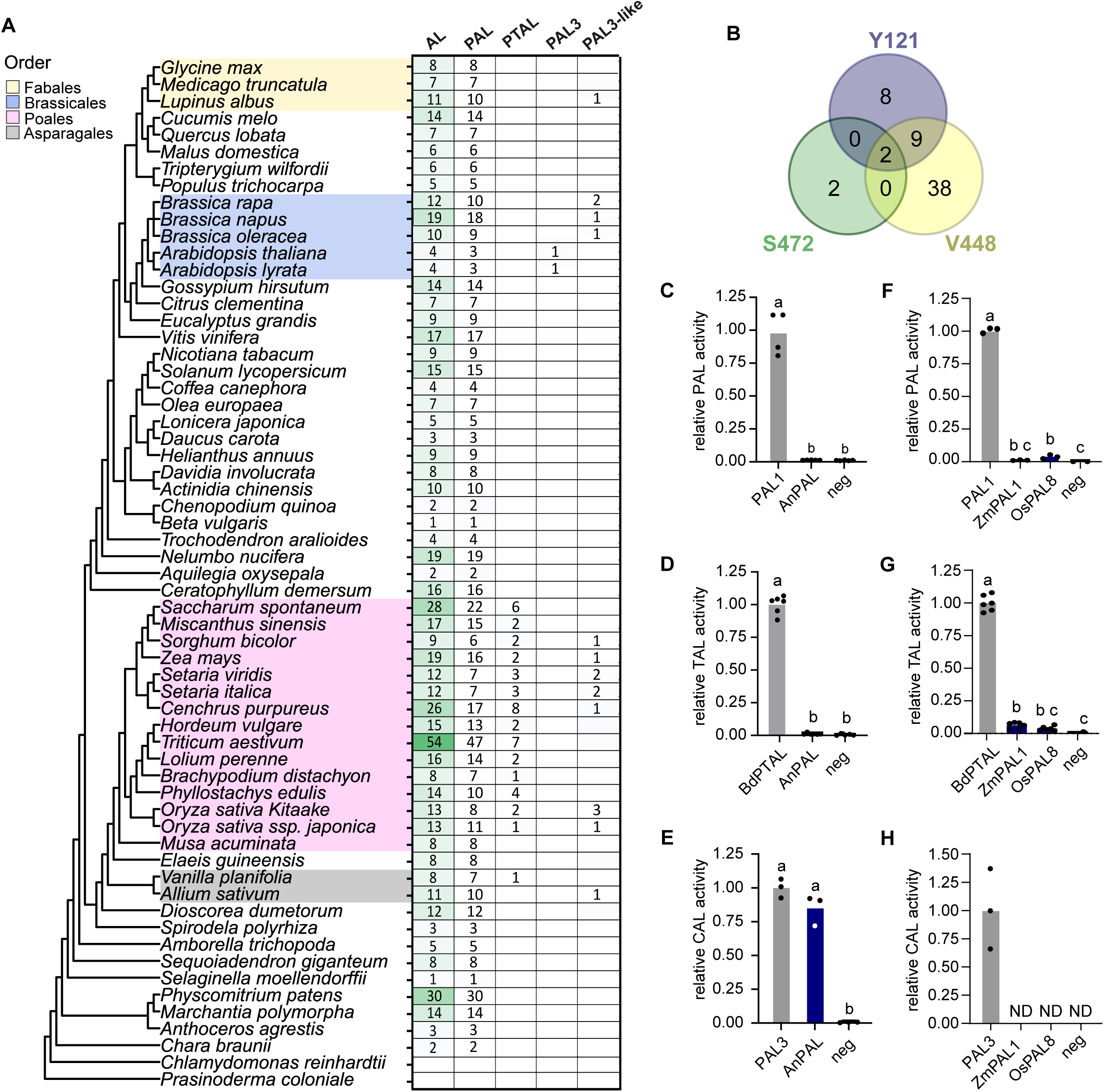
Distribution of ammonia lyases in plants. (**A**) Phylogenetic tree covering a broad range of plant species coupled to the number of predicted ammonia lyases in the genome sequences of the corresponding species. The ammonia lyase sequences are further categorized as PAL, PTAL, PAL3 or PAL3-like. The distinction between the different ammonia lyases was done based on the amino acids present at the substrate specificity site in the catalytic pocket as defined by Watts et al (2006) (i.e. phenylalanine for PAL and histidine for PTAL) and this study (tyrosine for PAL3). PAL3-like sequences share the tyrosine at the substrate specificity site with PAL3 (Y121), but lack S472 in the catalytic pocket crucial for CAL activity. (**B**) Venn diagram summarizing the number of ammonia lyases sharing at least one of the three amino acid residues related to CAL activity in PAL3 (i.e. Y121, V448 or S472). (**C-E**) Enzyme assays using heterologously produced PAL1 of Arabidopsis, AnPAL (PAL of *Arabis nemorensis)*. (**F-H**) Enzyme assays using heterologously produced OsPAL8 (PAL3-like from rice) and ZmPAL1 (PAL3-like from maize). In (**C** and **F**) L-phenylalanine was used as substrate and the absorbance at 270 nm was used as a proxy of cinnamic acid production. Absorbance values of the different samples were normalized to PAL1. In (**D** and **G**) L-tyrosine was used as substrate and the absorbance at 310 nm was used as a proxy of *p*-coumaric acid production. Absorbance values were normalized to the positive control *Brachypodium dystachion* PTAL. In (**E** and **H**) 3-carboxy-tyrosine was used as substrate and product formation was analysed using HPLC-MS. Values represented as product formation detected by MS/MS was normalized to the corresponding substrate amount detected by MS/MS and subsequently normalized to the conversion of the PAL3 positive control. AL: ammonia lyase, CAL: carboxy-tyrosine ammonia lyase, HPLC-MS: high performance liquid chromatography mass spectrometry, PAL: phenylalanine ammonia lyase, PTAL: phenylalanine tyrosine ammonia lyase.

Intriguingly, 17 putative ammonia lyases possessed a PAL3-specific tyrosine at their substrate specificity site (corresponding to Y121 in PAL3 of *A. thaliana*), but lacked the PAL3-key catalytic serine residue we identified (S472) (**Supplementary Fig. 12**). Together this indicates that these enzymes will have neither CAL nor any known ammonia lyase activity, suggesting that their native substrates remain to be identified. To verify this assumption, we cloned the CDS of such *PAL3*-like genes of rice (*OsPAL8*) and maize (*ZmPAL1*) and produced the corresponding enzyme in *E. coli*. In line with the prediction, these enzymes displayed no CAL activity, and no or only negligible PAL and TAL activity (**Fig. 4F-H**).

## DISCUSSION

Ammonia lyases are enzymes that catalyse the deamination of amino acids, resulting in the formation of their corresponding α,β-unsaturated carboxylic acids. The only members of this enzyme class in plants are PALs, which convert L-phenylalanine into cinnamic acid, and PTALs, which additionally accept L-tyrosine as a substrate to form *p*-coumaric acid^2^. These reactions represent the entry point into the phenylpropanoid pathway, a central metabolic route responsible for the biosynthesis of lignin, flavonoids, and a diverse array of other specialized metabolites essential for growth, development, and environmental adaptation^7^. Through their position at a major metabolic branch point, PALs and PTALs exert significant control over carbon flux into phenylpropanoid-derived compounds^10,13,20^.

Based on genomic annotation, PAL represents the only ammonia lyase identified in dicots like *Arabidopsis thaliana*, where it is encoded by a small gene family comprising four members. We demonstrated that PAL3 has been miscategorized for decades as it does not deaminate L-phenylalanine, but converts 3-carboxy-tyrosine into carboxy-*p*-coumaric acid. Based on this novel function we suggest to rename PAL3 to 3-carboxy-tyrosine ammonia lyase or CAL. This reclassification of PAL3 brings the number of true PAL enzymes in Arabidopsis to three, and adds a novel member of aromatic amino acid ammonia lyases to the existing categories of aromatic ammonia lyases acting on L-phenylalanine (PALs), L-tyrosine (TALs), and L-histidine (HALs). This finding clarifies why only negligible PAL or TAL activity was demonstrated for PAL3 using heterologously produced enzymes (Cochrane, Davin, and Lewis 2004), and why no difference in PAL activity was detected in crude enzyme extracts of *pal1/2* and *pal1/2/3* mutants, while a severe drop in PAL activity was observed in *pal1/2/4* and *pal1/2/3/4* compared to *pal1/2* mutants^10^. In line with these observations, we did not observe an additional negative effect on rosette size when *PAL3* was disrupted in the *pal1/2* background, while disrupting *PAL4* severely affected plant growth.

The substrate specificity of aromatic amino acid ammonia lyases was previously thought to be largely determined by a single, highly conserved amino acid residue located within the catalytic pocket^16^. In PALs, this position is occupied by a phenylalanine residue, which confers strict specificity toward L-phenylalanine. In contrast, PTALs possess a histidine residue at the corresponding position, enabling the accommodation and efficient conversion of both L-phenylalanine and L-tyrosine^16^.

Mutating the phenylalanine to a histidine in Arabidopsis PAL1 increased the enzymes activity towards L-tyrosine, while substituting the histidine to a phenylalanine in *Cereibacter sphaeroides* (previously known as *Rhodobacter sphaeroides*) resulted in a complete switch from TAL to PAL activity^16^. The novel identified CAL enzyme has a tyrosine at the substrate specificity site (i.e. Y121). Surprisingly, while this position severely impacted PAL and CAL activity, it was not the key substrate determinant in CAL. Compared to the catalytic pocket of PAL, two additional amino acid switches were identified, namely V448 and S472, of which only the second had a major impact on the substrate specificity. Based on the criterion that S472 represents a key residue required for CAL activity, this class of ammonia lyases has so far been identified in four species that are part of the Brassicaceae family. In this family the enzymes have evolved two times independently, once in the *Arabidopsis* genus and once in *A. nemorensis*.

In addition to identifying a novel type of ammonia lyase, our research uncovered the 3-carboxy-phenylpropanoids as a distinct, novel class of specialized metabolites within plant metabolism. Interestingly, the substrate 3-carboxy-tyrosine was already reported over fifty years ago as a shikimate-derived compound using radioactive labelling studies in *Reseda lutea*^21,22^. Recent metabolic profiling has confirmed these findings, demonstrating that 3-carboxy-tyrosine is derived from chorismate via isochorismate synthase (ICS), which converts chorismate to isochorismate (**Supplementary Fig. 13**)^23–25^. Besides being a precursor for 3-carboxy-phenylpropanoids, isochorismate acts as a branching point for the synthesis of the phytohormone salicylic acid and vitamin K’s (menaquinones, phylloquinones), which are involved in plant defence and photosynthesis, respectively^25^. Another enzyme, reversal of SAV3 phenotype (VAS1), acts in this metabolic side branch and was identified as the enzyme involved in the synthesis of 3-carboxy-tyrosine and 3-carboxy-phenylalanine from 3-carboxy-hydroxyphenylpyruvate and 3-carboxy-phenylpyruvate, respectively^23^. The addition of CAL and the downstream 3-carboxy-phenylpropanoids adds another piece to the complex puzzle of the plants metabolic network.

The physiological role of this newly characterised enzyme and novel class of associated specialized metabolites remains poorly understood. One hypothesis supported by several studies is that CAL and the carboxy-phenylpropanoids might be implicated in pathogen defence. For example, knocking-out *CAL* in a *pal1/2/4* mutant background leads to a notable reduction in both basal and *Pseudomonas syringae*-induced accumulation of salicylic acid^10^. Furthermore, the phytotoxin coronatine, which enhances the pathogenic effect of *Pseudomonas syringae* in plants, has been shown to supress *CAL* expression in Arabidopsis^26^. Additionally, Guo et al. (2016) showed that the *CAL* promoter is activated by the defence-related transcription factor *ethylene response factor 1 (ERF1)-like* from cotton (*Gossypium barbadense*) upon overexpression in Arabidopsis^27^. In the context of pathogen-resistance, carboxy-phenylpropanoids may serve as signalling molecules or as pathogen-targeting toxins^27^. To fully elucidate the role of *CAL* in biotic stress responses, comprehensive phenotypic analysis should be conducted, assessing the performance of *CAL* loss- and gain-off function mutants under various biotic conditions.

In conclusion, our research provides novel insights into the functional diversity of ammonia lyases through the identification of CAL as a novel enzyme distinct from PALs and (P)TALs. This discovery not only increases the functional diversity of ammonia lyases, it also introduces the 3-carboxy-phenylpropanoids as a novel class of specialized metabolites. Besides providing novel insights into previously unexplored aspects of plant metabolism, it opens new avenues for biotechnological and industrial applications.

## LIST OF AUTHOR CONTRIBUTIONS

CVB, AVdV, RV and BV designed the experiments; CVB, AVdV and BV performed the experiments; JG analysed the PAL3 enzymatic samples by HPLC-MS and synthesized 3-carboxy-tyrosine; SV processed the metabolomics UPLC-MS samples; RV and SD analysed the metabolomics data; CX and BV performed the protein purification experiments; FAS performed the phylogenetic analysis; HM and TB performed structural analyses and docking; CVB, AVdV and BV wrote the manuscript; JVdE, BD, UB, and WB discussed the project and complemented the writing together with HM and TB.

## ACKNOWLEDGEMENTS

We would like to extend our gratitude to Dr Meng Peng for the assistance with protein purification, Dr Ward Decaestecker for the assistance designing the CRISPR/Cas9 constructs and Dr Willem Desmedt for the *pPAL3::GUS* reporter lines.

## FUNDING

This work was funded by Bijzonder Onderzoeksfonds bof/baf/4y/2024/01/773 (CVB) and The China Scholarship Council DOCT/005782 (CX)

## Legends to supplementary figures

**Supplementary Figure 1: Overview of the mutations in the CRISPR/Cas9-mediated *pal* mutant lines.** Protospacer and PAM (underlined) sequences are represented in 5’ to 3’ direction and are situated on the sense strand, except for the second protospacer and PAM of *PAL4* which are situated on the antisense strand. The red arrows indicate the positions of the guide RNAs in the different *PAL* genes. CRISPR/Cas9-mediated edits in the sequences are represented in red. bp: base pair, CRISPR: Clustered Regularly Interspaced Short Palindromic Repeats, PAL: phenylalanine ammonia lyase, PAM: protospacer adjacent motif, WT: wildtype.

**Supplementary Figure 2: Rosette phenotypes of *pal* single mutants 21 DAS.** Plants were grown horizontally on ½ MS medium under long day growth conditions (16 h light / 8 h dark) at 21 °C. Scale bar represents 1 cm. DAS: days after stratification.

**Supplementary Figure 3: SDS-PAGE of purified PAL3 protein.** PAL3 was heterologously expressed as a His6x-MBP fusion protein. Fraction 1 and 2 correspond to proteins eluted with 50 or 100 mM imidazole, respectively. Mar: marker, MBP: maltose binding protein, Lys: crude lysate, PAL: phenylalanine ammonia lyase, Sup: supernatant, kDa: kilodalton.

**Supplementary Figure 4: Multiple alignment of Arabidopsis *PAL* genes.** Amino acid residues of the catalytic pocket are indicated in bold. Black residues represent amino acids that are conserved between PAL1, 2 and 4, and PAL3. Blue residues represent amino acids that are different between PAL1, 2 and 4. MIO group is indicated in pink. MIO: 4-methylideneimidazol-5-one, PAL: phenylalanine ammonia lyase

**Supplementary Figure 5: Enzyme assay using all 20 amino acids as substrates**. Amino acids were analysed via GC-MS. Samples were normalized based on total peak area for all detected amino acids. Statistical significance was determined using a Students t-test (*p* < 0.01). n = 4. GC-MS: gas chromatography-mass spectrometry, HAL: histidine ammonia lyase, PAL: phenylalanine ammonia lyase,

**Supplementary Figure 6: Expression pattern of *PAL3*. (A**) *in silico* expression pattern of *PAL3* in different tissues of *A. thaliana* over different life stages reported on the eFP Browser 2.0. (**B-D**) *PAL3* promotor activity analysis using GUS reporter lines 7 DAS. (**E-H**) *PAL3* promotor activity analysis using GUS reporter lines 14 DAS. Plants were grown horizontal plates on ½ MS medium under long day growth conditions (16 h light / 8 h dark) at 21 °C. Scale bar in C, D, G and H represents 2 mm, and scale bar in B, E and F represents 0.2 mm. DAS: days after stratification

**Supplemental Figure 7. Structural elucidation of metabolites via MS/MS spectra**

**Supplementary Figure 8: Substrate inhibition of PAL3 and PAL1 with their respective substrates.** (**A**) *In vitro* enzyme assays using heterologously produced PAL3 and 3-carboxy-tyrosine as substrate at concentrations between 0.15 and 1.25 mM. The PAL3 product 3-carboxy-*p*-coumaric was measured by HPLC-MS. (**B**) *In vitro* enzyme assay using heterologously produced PAL1 and L-phenylalanine as substrate at concentrations between 0.15 and 1.5 mM. Absorbance at 270 nm was used as a proxy of cinnamic acid production. HPLC-MS: high performance liquid chromatography mass spectrometry, PAL: phenylalanine ammonia lyase.

**Supplementary Figure 9: Phylogenetic tree of ammonia lyases in plants.** The left panel shows a phylogenetic tree covering a broad range of plant species. The middle panel shows the number of predicted ammonia lyases in each of the plant species depicted in the phylogenetic tree. The right panel shows the number of different ammonia lyases. A distinction was made between PAL, PTAL, PAL3 (with CAL activity demonstrated in this paper) and PAL3-likes. The distinction between the different ammonia lyases was done based on the amino acids present at the substrate specificity site in the catalytic pocket as defined by Watts et al (2006) (i.e. phenylalanine for PAL and histidine for PTAL) and this study (tyrosine for PAL3). PAL3-like sequences share the tyrosine at the substrate specificity site with PAL3, but lack the serine in the catalytic pocket crucial for CAL activity. An empty cell indicates no sequences fulfil the defined criteria. AL: ammonia lyase, CAL: 3-carboxy-tyrosine ammonia lyase, PAL: phenylalanine ammonia lyase, PTAL: phenylalanine tyrosine ammonia lyase.

**Supplementary Figure 10: Analysis of homologous PAL3 sequences possessing the PAL3 triad. (A)** Multiple alignment of homologous PAL3 sequences possessing the PAL3 triad in their catalytic pocket. The highlighted positions refer to the positions of the amino acids in PAL3. Sequences were obtained via Uniprot. (**B**) Sequence similarity (lower left) and identity (upper right) for the homologous PAL3 sequences introduced in (A). Values are represented as percentages. Blosum45 was used with threshold 0 and data analyses was done in Geneious. PAL: phenylalanine ammonia lyase

**Supplementary Figure 11: Phylogenetic tree of ammonia lyases in the Brassicaceae.** Dark blue cell indicates at least one ammonia lyase with predicted CAL activity based on presence of all three amino acid switches (Y121, V448 and S472 in Arabidopsis PAL3). Light blue cell indicates no ammonia lyases with predicted CAL activity due to the lack of these three amino acid switches. AL: e ammonia lyase.

**Supplementary Figure 12: Multiple sequence alignment of the ammonia lyases sharing at least one of the three amino acid residues related to CAL activity in PAL3 (i.e. Y121, V448 or S472)**. CAL: 3-carboxy-tyrosine ammonia lyase, PAL phenylalanine ammonia lyase.

**Supplementary Figure 13: Biosynthesis pathways to the aromatic amino acids, phenylpropanoids, 3-carboxy-amino acids and 3-carboxy-phenylpropanoids.** ADH: arogenate dehydrogenase, ADT: arogenate dehydratase, CM: chorismate mutase, C4H: cinnamate-4-hydroxylase, ICS: isochorismate synthase, PAL: phenylalanine ammonia lyase, PTAL: phenylalanine tyrosine ammonia lyase PPA-AT: prephenate aminotransferase, TAL: tyrosine ammonia lyase, VAS1: reversal of *sav3* phenotype.

**Supplementary Figure 14: Synthesis of 3-carboxy-tyrosine.**

## Legend to supplementary table

**Supplementary Table 1**: **Sequences of oligonucleotides used in this work**. Primer sequences are represented in 5’-3’ direction using the coding strand as reference. FP, forward primer; PAL, phenylalanine ammonia lyase; PTAL, phenylalanine tyrosine ammonia lyase; RP, reverse primer.

## MATERIALS AND METHODS

### Plant material and growth conditions

All *Arabidopsis thaliana* lines described in this study are in the Columbia-0 (Col-0) ecotype background. When grown *in vitro*, seeds were surface sterilized using vapor-phase seed sterilization. Briefly, dry seeds were placed in open 1.5-mL microcentrifuge tubes and transferred to a desiccator. Chlorine gas was generated by combining 100 mL bleach (12-13 % sodium hypochlorite solution; Chem-Lab) with 3.5 mL concentrated hydrochloric acid (37 %; Supelco) in a separate open vessel within the desiccator, which was immediately sealed to allow exposure of the seeds to chlorine vapor. After overnight incubation at room temperature, the desiccator was opened in a fume hood and seeds were allowed to off-gas briefly. Sterilized seeds were transferred to round (diameter 14.5 cm; Greiner Bio-One) or square (12 x 12 cm; Corning) plates containing 0.8 % (w/v) agar-solidified 1/2 Murashige and Skoog (MS) medium containing per liter 1.5 g Murashige and Skoog basal salt mixture powder (Duchefa), 7.14 g saccharose (Chem-Lab), 0.36 g 2-(N-morpholino)ethanesulfonic acid (MES) monohydrate (Duchefa), and 8 g plant tissue culture grade agar No. 4 (Neogen) (pH 5.7; adjusted with 1 M KOH). After sowing, plates were incubated in the dark at 4 °C for three days and afterwards transferred to long-day growth conditions (16 h light / 8 h dark) at 21 °C and incubated in either vertical or horizontal orientation. Light was provided by full-spectrum Osram L 58W/840 Lumilux Cool White fluorescent lamps (FL), delivering a photosynthetic photon flux density (PPFD) of 90 µmol/m².s at shelf level, measured using a SpectroSense2+ quantum sensor (Skye Instruments Ltd, UK). When grown in soil, dry Arabidopsis seeds were sown directly onto the surface of pre-soaked peat-based Jiffy pellets, which were maintained in trays. Plants were grown at 21 °C in a controlled-environment growth chamber under long-day conditions (16 h light / 8 h dark) at 21 °C.

### Plant phenotyping

The quantification of the projected rosette area was performed on Arabidopsis plants grown horizontally on 1/2 MS medium in round petri dishes (diameter 14.5 cm; Greiner Bio-One). At 21 days after stratification (DAS), plates were digitized using an Expression 11000XL flatbed scanner (Epson) operated with SilverFast 8 software, and rosette surface area was measured using ImageJ software (Schneider et al. 2012). Images were converted to binary (black-and-white), after which the surface area of each rosette was measured. The results were processed using GraphPad Prism10 software.

### Synthesis of 3-carboxy-tyrosine (Supplementary Fig 14)

#### Synthesis of Boc-3-iodoTyrOMe

(Leroux et al. 2019): BocTyrOMe (5.00 g, 1.0 equiv; Merck Life Science BV, Belgium) and sodium iodide (3.05 g, 1.2 equiv) were dissolved in dimethylformamide (DMF; 80 mL) in a round bottom flask equipped with a magnetic stirring bar and septum. A solution of Chloramine-T trihydrate (5.72 g, 1.2 equiv) in DMF (40 mL) was added to the flask over 2.5 h at -10 °C under argon atmosphere. Afterwards the reaction mixture was kept overnight at room temperature. After quenching with 400 mL water, the product was extracted with 4x 500 mL ethylacetate followed by washing with 3x 200 mL brine. The crude extract was redissolved in 250 mL diethyl ether and washed 6x with saturated ammonium chloride in water, followed by drying over magnesium sulphate. The crude product was purified by flash column chromatography (dichloromethane/acetone 97:3) yielding 4.18 g pure Boc-3-iodoTyrOMe. **^1^H-NMR (400 MHz, CDCl_3_)**: δ [ppm] = 7.42 (s, 1H), 6.97 (d, J=8.2, 1H), 6.83 (d, J=8.2, 1H), 6.09 (s, 1H), 5.06 (d, J=7.8, 1H), 4.50 (m, 1H), 3.71 (s, 3H), 3.03 (dd, J=13.9, 6.0, 1H), 2.91 (dd, J=13.9, 6.0, 1H), 1.42 (s, 9H). **^13^C-NMR (100 MHz, CDCl_3_)**: δ [ppm] = 172.2, 155.1, 154.2, 139.0, 130.8, 129.8, 115.0, 85.2, 80.2, 54.4, 52.3, 36.9, 28.3. **HRMS (ESI)**: m/z calc. for [M+Na^+^; C_15_H_20_INO_5_Na^+^]: 444.0278; found: 444.0299. **IR (Diamond-ATR, neat)**: cm^-1^ = 3358 (w), 2976 (w), 1681 (s), 1604 (w), 1573(w), 1503 (m), 1438 (w), 1414 (w), 1392 (w), 1366 (m), 1289 (w), 1250 (w), 1216 (m), 1158 (s), 1058 (m), 1036(w), 1016 (m), 823 (w), 806 (w), 742 (w), 665 (w).

#### Synthesis of Boc-3-(methoxycarbonyl)TyrOMe

(Lokolkar and Bhanage 2023): To a 250-mL stainless-steel pressure reactor, Boc-3-iodoTyrOMe (3.53 g, 1.0 equiv), *tetrakis*(triphenylphenylphosphine)palladium[0] catalyst (485 mg, 0.05 equiv), diisopropylethylamine (3.64 mL, 2.5 equiv) and 170 mL methanol were added. The reactor was pressurized with 7.5 bar CO gas and heated to 85 °C for 3 h. Then, the reaction mixture was concentrated in vacuo, 250 mL water was added, followed by extraction with 4x 250 mL diethyl ether. The extract was washed with 3x 100 mL brine, followed by drying over magnesium sulphate. The crude product was purified by flash column chromatography (*n*-hexane/acetone 85:15) yielding 2.63 g pure Boc-3-(methoxycarbonyl)TyrOMe. **^1^H-NMR (400 MHz, aceton-d_6_)**: δ [ppm] = 10.60 (s, 1H), 7.70 (s, 1H), 7.41 (d, J=8.2, 1H), 6.89 (d, J=8.2, 1H), 6.17 (d, J=7.7, 1H), 4.35 (m, 1H), 3.94 (s, 3H), 3.66 (s, 3H), 3.08 (dd, J=13.9, 5.2, 1H), 2.92 (dd, J=13.9, 8.8, 1H), 1.33 (s, 9H). **^13^C-NMR (100 MHz, aceton-d6)**: δ [ppm] = 173.6, 171.8, 161.8, 156.7, 138.3, 131.8, 129.7, 118.7, 113.4, 79.9, 56.5, 53.4, 52.8, 37.8, 29.0. **HRMS (ESI)**: m/z calc. for [M+Na^+^; C_17_H_23_NO_7_Na^+^]: 376,1372; found: 376.1373. **IR (Diamond-ATR, neat)**: cm^-1^ = 3384 (w), 2957 (w), 1732 (m), 1672 (s), 1614 (w), 1596 (w), 1512 (m), 1488 (m), 1442 (m), 1394 (w), 1366 (w), 1341 (w), 1283 (m), 1249 (s), 1217 (s), 1160 (s), 1093 (m), 1052 (w), 1010 (m), 976 (w), 928 (w), 917 (w), 850 (w), 835 (w), 798 (m), 775 (w), 742 (m), 677 (w). **CHIRAL HPLC**: Daicel Chiralcel ODH 4,6 x 250 mm, *n*-hexane/ehtanol 97:3, 1 mL / min, 35 °C: shows only one peak at 11.8 min.

#### Synthesis of 3-carboxy-L-tyrosine hydrochloride

(Arnold 1977; Arnold 1985): Boc-3-(methoxycarbonyl)TyrOMe (2.49 g, 1 equiv) was dissolved in 4 M HCl (70.5 mL, 30 equiv) in a round bottom flask equipped with a magnetic stirring bar, reflux condenser and argon balloon. The flask was heated to 100 °C for 3.5 h. The hot mixture was filtered over a glass filter and the filtrate evaporated to dryness. The whole procedure was repeated a second time, after which 20 mL toluene was added. The mixture was concentrated in vacuo, yielding 1.30 g white powder. Analytical RP-HPLC indicated a purity exceeding 99 % (214 nm). **^1^H-NMR (400 MHz, DMSO-d_6_)**: δ [ppm] = 8.47 (s, 3H), 7.69 (s, 1H), 7.42 (d, J=8.5, 1H), 6.91 (d, J=8.5, 1H), 4.08 (s, 1H), 3.09 (d, J=6.2, 1H). **^13^C-NMR (100 MHz, DMSO-d_6_)**: δ [ppm] = 171.8, 170.3, 160.4, 136.9, 131.3, 125.5, 117.3, 112.9, 53.2, 34.6 **HRMS (ESI)**: m/z calc. for [M+H^+^; C_10_H_12_NO_5_^+^]: 226.0710; found: 226.0699. **IR (Diamond-ATR, neat)**: cm^-1^ = 2967 (m), 2903 (m), 2608 (w), 2557 (w), 1738 (m), 1664 (s), 1616 (w), 1589 (w), 1488 (m), 1437 (s), 1360 (w), 1287 (w), 1208 (s), 1142 (w), 1128 (w), 1089 (w), 1061 (w), 963 (w), 882 (w), 848 (w), 825 (w), 795 (m), 766 (w), 733 (w), 690 (m), 670 (m), 609 (w). **Optical Rotation**: [α]_D_^25^ : -19,6°. **Melting Point**: 219 °C (decomposition).

### Generation of *pal* CRISPR lines

To generate *pal1, pal2, pal3*, *pal4*, *pal1/2*, *pal1/2/3*, *pal1/2/4* and *pal1/2/3/4* CRISPR/Cas9-mediated mutant lines, gRNAs targeting the Arabidopsis *PAL1 (*AT2G37040 ), *PAL2 (*AT3G53260), PAL*3 (*AT5G04230), and *PAL4 (*AT3G10340) genes were designed (**Supplementary Fig 1; Supplementary Table S1**, primers 1-6) using the online tools CRISPR-P 2.0 (http://crispr.hzau.edu.cn/CRISPR2/) and CRISPR-PLANT v2 (https://crispr-plantv2.bastianminkenberg.com/). For both *PAL1* and *PAL2*, a single gRNA was designed and the protospacer sequence of each gRNA was ordered as a sense (ATTG-N20, with N20 being the protospacer of the gRNA) and an antisense (AAAC-N20, with N20 being the reverse complement of the protospacer of the gRNA) oligonucleotide. Complementary oligonucleotides (100 µM stocks) were annealed by combining 1 µL of each oligonucleotide with 48 µL of Milli-Q (MQ) water. The mixture was subjected to thermal denaturation and annealing using a thermocycler (Bio-Rad Model T100), with the following program: 95 °C for 5 min, followed by 85 °C for 30 s and 25 °C for 30 s. For *PAL3* and *PAL4*, two gRNAs were used and the protospacer sequences of each pair of gRNAs was ordered as a long forward oligonucleotide (TTTTGGTCTCAATTG-N20-GTTTTAGAGCTAGAAATAGC, with N20 being the protospacer of gRNA1) and a long reverse oligonucleotide (TTTTGGTCTCAAAAC-N20-CAATCACTACTTCGACTC, with N20 being the reverse complement protospacer of gRNA2) primer. For all four *PAL* genes, the protospacer sequences of the gRNAs (annealed or PCR product) was verified by agarose gel electrophoresis and gel-purified using the GeneJet Gel Extraction Kit (Thermo Fisher Scientific). Subsequently, the protospacer sequences of the four PAL genes were cloned in pGG-A-AtU6-ccdB-B, pGG-B-AtU6-ccdB-C, pGG-C-AtU6-ccdB-D, and pGG-D-AtU6-ccdB-D Golden Gate donor vectors, respectively, to generate the gRNA entry clones. Recombinant clones were selected in *E. coli* DH5α cells using carbenicillin (100 µg/mL) as a selection marker. Verified gRNA entry clones were recombined with a Golden Gate-compatible destination vector pFASTRK-AtCas9-A-ccdB-G (9_43) to generate the CRISPR expression clones. Recombinant clones were selected in *E. coli* DH5α cells using spectinomycin (50 µg/mL) as a selection marker. Successful cloning of both entry and expression clones was confirmed by Sanger sequencing (Eurofins Scientific, Belgium) (**Supplementary Table S1,** primers 7-12).

Sequence-verified expression clones were introduced into *Agrobacterium tumefaciens* strain C58C1 by electroporation using a MicroPulser electroporator (Bio-Rad). Following transformation, cells were allowed to recover under non-selective conditions for 2 h at 28 °C and 180 rpm in 1 mL of liquid Yeast Extract Broth (YEB; 1 g/L beef extract, 5 g/L yeast extract, 5 g/L peptone, 5 g/L sucrose, 0.5 g/L magnesium sulphate; pH 7.0). After the recovery period, cultures were plated onto YEB agar supplemented with rifampicin (100 mg/L), gentamicin (25 mg/L), and spectinomycin (100 mg/L) to select for successfully transformed Agrobacterium cells. Antibiotic-resistant colonies were subsequently screened by colony PCR.

The transformation of Arabidopsis was done by floral dipping. Briefly, Agrobacterium strains carrying the binary vector of interest were grown overnight at 28 °C and 200 rpm in 5 mL YEB supplemented with rifampicin, gentamicin, and spectinomycin as before. This preculture was used to inoculate 200 mL YEB, which was grown overnight under similar growth conditions. The next day, cells were harvested by centrifugation (21,000 x g for 15 min), and the pellet was resuspended in infiltration medium consisting of 100 g/L (w/v) saccharose and 0.02 % (v/v) Silwet L-77. Flowering Arabidopsis plants were inverted and submerged in the Agrobacterium suspension for approximately 30 s with gentle agitation to ensure thorough wetting of floral tissues. Following dipping, plants were covered with a transparent lid or plastic dome to maintain high humidity for 24 h, after which they were returned to standard growth conditions. Seeds (T1 generation) were collected approximately four weeks after dipping, and screened under fluorescent light, using an epifluorescence microscope (Leica DFC 7000T; excitation and emission maxima at 558 nm and 605 nm, respectively) to select transgenic lines. To obtain transgene-free, biallelic edited *pal* lines, fluorescent-free seeds were selected over three subsequent generations.

During the selection, plant editing was confirmed by genotyping using genomic DNA extracted according to the Edwards method. Briefly, one fresh leaf was collected from individual seedlings (10 DAS) and transferred to an empty 8-strip 1.1-mL deep well microtube (National Scientific Supply Co, Inc.). To each tube, two stainless steel beads (4 mm diameter) were added, whereupon the tubes were closed with caps. Twelve strips were arranged in a microrack to match a standard 96-well microplate footprint. During sampling tubes were chilled on dry ice. Following collection, samples were immediately snap-frozen in liquid nitrogen and stored at −80 °C until further processing. Frozen samples were homogenized using a Retsch MM 400 Mixer Mill equipped with a 96-well adapter set pre-cooled to −80 °C. Homogenization was performed at 20 Hz for 30 s. Subsequently, 400 µL of preheated (60 °C) Edwards extraction buffer (200 mM Tris-HCl, 250 mM NaCl, 25 mM EDTA, 0.5 % SDS; pH 7.5) was added to each tube. Samples were mixed thoroughly by inversion and incubated at 60 °C for 10 min. Cellular debris was removed by centrifugation at 13,000 × g for 15 min at room temperature using an Eppendorf 5810R centrifuge. Following centrifugation, 300 µL of the supernatant was transferred to a fresh tube, and genomic DNA was precipitated by adding an equal volume of isopropanol. After centrifugation at 13,000 × g for 20 min, the resulting DNA pellet was washed with 100 µL of 70 % (v/v) ethanol, air-dried, and resuspended in 100 µL of sterile MQ water. Genotyping was performed by PCR, using 3 µL of crude genomic DNA as templated in a 50 µL total reaction volume, containing 1x Green Flexi GoTaq Reaction Buffer, 0.1 mM of each dNTP, 0.3 µM of each primer (**Supplementary Table S1,** primers 15-31), 0.625 mM MgCl₂, and 1 U Taq DNA polymerase (Promega). PCR amplification was carried out under standard cycling conditions: initial denaturation at 95 °C for 5 min; 36 cycles of 95 °C for 30 s, 54 °C for 30 s, and 72 °C for 1 min per kb; followed by a final extension at 72 °C for 5 min. PCR products were resolved on a 1.5 % (w/v) agarose gel, and positive samples were further purified using HighPrep PCR (MagBio) according to the manufacturer’s instructions. The purified DNA was used for Sanger sequencing (Eurofins Scientific, Belgium). For each mutant, two independent transgene-free, biallelic edited lines were generated.

### Generation of GUS reporter lines

Genomic DNA was isolated from rosette leaves of *in vitro* grown *A. thaliana* plants at 21 DAS, using the Wizard^®^ Genomic DNA Purification Kit (Promega) according to the manufacturer’s instructions. The quality of the purified DNA was assessed by agarose gel electrophoresis. The genomic DNA was diluted 1:1000 in sterile water, and 1 µL was used as template to amplify a promoter fragment of 2235 bp upstream of the annotated transcriptional start site of the *PAL3*-coding sequence using a nested PCR approach. In the first PCR, the target region was amplified using an outer primer pair (**Supplementary Table S1,** primers 32 & 33 under following PCR conditions: 98 °C for 30 s; 30 cycles of 98 °C for 15 s, 60 °C for 25 s, and 72 °C for 1.5 min; followed by a final extension at 72 °C for 5 min. An aliquot of the primary reaction product was 1:100-diluted in water and 1 µL was subsequently used as template in a second amplification round with an inner (nested) primer pair (**Supplementary Table S1**, primers 34 & 35) specific to a sequence internal to the first amplicon. The inner primers contained *attB*recombination sequences at their 5′ ends, allowing downstream Gateway cloning. PCR conditions were as follows: 98 °C for 30 s; 36 cycles of 98 °C for 15 s, 60 °C for 25 s, and 72 °C for 1.5 min; followed by a final extension at 72 °C for 5 min. Both PCRs were carried out using Q5^®^ High-Fidelity DNA Polymerase (New England Biolabs) according to the manufacturer’s instructions. The amplified fragment was verified by agarose gel electrophoresis and gel-purified using the GeneJet Gel Extraction Kit (ThermoFisher Scientific) prior to recombination into a Gateway donor vector *pDONR221* via BP recombination to generate the promoter entry clone. Recombinant clones were selected in *E. coli* DH5α cells using kanamycin (50 mg/L) as selection marker. Correct insertion of the promoter fragment was verified by colony PCR using *attB1* and *attB2* primers (**Supplementary Table S1,** primers 13 & 14). Sequence integrity was subsequently confirmed by Sanger sequencing. The verified promoter entry clone was recombined with a Gateway-compatible destination vector *pKGWFS7* containing a promoterless *GUS* reporter gene using LR recombination (*pPAL3*). The resulting expression clone placed the *PAL3* promoter fragment upstream of the reporter coding sequence. Positive clones were initially screened by colony PCR, using the same primer pair targeting the recombination-flanking regions as described before. The integrity of the insert was subsequently verified by Sanger sequencing across the promoter–reporter junction. The final binary construct was introduced into *A. tumefaciens* and subsequently transformed into *A. thaliana* plants via floral dipping as explained in the section on the generation of *pal* CRISPR lines above. Transgenic lines were selected on 1/2 MS medium containing 50 mg/L kanamycin. The reporter gene expression was analysed in T1 and subsequent generations by GUS analysis as described below.

### GUS expression analysis

For the expression analysis in seedlings, seeds of two independent *PAL3* reporter lines were sown on 1/2 MS medium and placed vertically in the growth room under long day growth conditions (16 h light, 8 h dark photoperiod). At 7 and 14 DAS, seedlings were harvested and immersed in 90 % (v/v) acetone pre-cooled to −20 °C, followed by incubation at 4 °C for at least 1 h. Subsequently, the acetone was carefully removed and replaced with NT buffer (100 mM Tris-HCl, 50 mM NaCl; pH 7.0) to eliminate residual solvent. Ferricyanide stock solution was prepared by dissolving 0.32 g K_3_[Fe(CN)_6_] (Sigma; CAS No. 13746-66-2) in 10 mL of NT-buffer. A ferricyanide working solution was then made by mixing 49 mL of NT-buffer with 1 mL of the ferricyanide stock. The plants were incubated in the ferricyanide working solution at 37 °C for at least 30 min. The final assay solution was prepared by mixing 24.5 mL of the ferricyanide working solution with 0.5 mL of the X-Gluc stock solution (0.0261 g X-Gluc (X-Gluc DIRECT) in 500 µL dimethyl sulfoxide). Finally, the ferricyanide working solution was replaced with the assay solution, and further incubated for at 37 °C until the desired staining was achieved. The staining reaction was stopped by replacing the assay solution with NT buffer. For microscopic analysis, seedlings or plant explants were carefully transferred onto glass microscope slides. A drop of lactic acid was applied to the tissue to facilitate destaining and improve optical clarity. Samples were then covered with a coverslip, ensuring even distribution of the clearing solution and minimizing air bubbles. Imaging was performed using an Olympus BX53 light microscope equipped with an UPlanFL N 20× objective lens (numerical aperture, NA = 0.5). All images were acquired under identical illumination and exposure settings to allow for consistent comparison between samples.

### Generating constructs for protein production

The *PAL1* coding sequenc (CDS) was PCR amplified using Q5^®^ High-Fidelity Polymerase (New England Biolabs) from an existing entry clone, with primers suitable for Gateway Cloning (*attB1* and *attB2*; **Supplementary Table S1,** primers 42 & 43). For *PAL3*, each of the three exons was amplified from the Arabidopsis genomic DNA that was extracted to clone *pPAL3*. The exons were amplified via nested PCR using Q5^®^ High-Fidelity Polymerase as described above with gene specific anchor primers (**Supplementary Table S1,** primers 36-41), allowing exon fusion. The amplified fragments were verified by agarose gel electrophoresis and gel-purified using the GeneJet Gel Extraction Kit (ThermoFisher Scientific). After purification, 5 µL of the PCR product was utilized to fuse exon 1 to exon 2, and exon 2 to exon 3, through another PCR with Q5^®^ polymerase. This PCR product was again gel-purified, and 5 µL of the purified product was used to fuse the exon 1-2 product to the exon 2-3 product. The final PCR products of *PAL1* and *PAL3* were gel-purified and used for Gateway cloning. Gateway recombination reactions were performed according to the manufacturer’s instructions (Invitrogen). For the BP reaction, the PCR-amplified *PAL1*and *PAL3* fragments flanked by *attB* sites were recombined with the donor vector *pDONR221* using BP Clonase™ II enzyme mix to generate the construct *pDONR221*-*PAL1* and *pDONR221*-*PAL3*, respectively. The reactions were incubated at 25 °C for 1 h and terminated by the addition of proteinase K, followed by transformation into heat shock competent *E. coli* DH5α cells by heat shock as described before. The selection of entry clones was done on lysogeny broth (LB) medium, supplemented with kanamycin (25 mg/L). For the LR reaction, the sequence verified entry clone was recombined with the *pDEST*-*HisMBP* destination vector, using LR Clonase™ II enzyme mix to generate the *pDEST-HisMBP-PAL1* and *pDEST*-*HisMBP*-*PAL3* expression constructs, respectively. The recombination reaction was incubated at 25 °C for 1 h, treated with proteinase, and transformed into competent *E. coli* DH5α for plasmid propagation and subsequent sequence verification. Transformed *E. coli* cells were grown on LB medium, supplemented with carbenicillin (100 µg/mL). Positive colonies were confirmed by colony PCR, using *attB1* and *attB2* primers (**Supplementary Table S1**).

Site-directed mutagenesis of PAL1 and PAL3 was performed by PCR, using primers containing the desired mutation (**Supplementary Table S1,** primers 44-57). After amplification and assembling the full length CDS using Q5^®^ High-Fidelity Polymerase, the amplicon was cloned in the *pDONR221* entry clone, and finally the *pDEST*-*HisMBP* expression vector as described above. The following mutations were introduced in PAL1: F144Y, I469V, and E493S. The following mutations were introduced in PAL3: Y121F, V448I, and S472E.

The CDS of *PALs* from rice (*OsPAL8*; LOC_Os11g48110) and maize (*ZmPAL1*; Zm00001d033286) were amplified from genomic DNA using the exon-fusion strategy previously applied for Arabidopsis *PAL3* (**Supplementary Table S1,** primers 58-71). Genomic DNA was extracted from leaf explants of *Zea mays* B104 and *Oryza sativa* L. subsp. *japonica* cv. ‘Nipponbare’ seedlings using the Wizard® Genomic DNA Purification Kit (Promega) according to the manufacturer’s instructions. The resulting CDS fragments were cloned into the *pDONR221* entry vector using the BP Clonase™ II enzyme mix to generate the construct *pDONR221-ZmPAL1* and *pDONR221-OsPAL8*, respectively. Positive clones were selected by colony PCR. For the LR reaction, the sequence verified entry clone was recombined with the *pDEST-HisMBP* destination vector using LR Clonase™ II enzyme mix to generate the *pDEST-HisMBP-ZmPAL1* and *pDEST-HisMBP-OsPAL8* expression constructs, respectively.

The *Arabis nemorensis PAL* (*AnPAL*) CDS was reverse translated from the Uniprot protein sequence, and ordered in the *pTWIST* entry vector from Twist Bioscience. The CDS of *Brachypodium distachyon* PTAL (*BdPTAL;* XP_003575396) and *Arabis nemorens*is PAL (AnPAL) were PCR amplified, using Q5^®^ High-Fidelity Polymerase (New England Biolabs) with primers suitable for Gateway Cloning (**Supplementary Table S1,** primers 72-75). After gel purification, the amplicons were cloned in the *pDONR221* entry clone, and finally *pDEST*-*HisMBP* expression vectors as described above.

### Heterologous protein production and purification

For protein production, chemical competent *E. coli* BL21 cells were transformed with the desired constructs by heat-shock as before. Transformants were selected on LB agar supplemented with carbenicillin (100 µg/mL). Following overnight incubation at 37 °C, a single colony was used to inoculate 5 mL of LB medium containing the same antibiotic and cultured overnight at 37 °C. Subsequently, 4 mL of this preculture was transferred into a 2-L Erlenmeyer flask containing 200 mL of LB medium supplemented with carbenicillin (100 µg/mL). The culture was grown at 37 °C with continuous shaking at 200 rpm until reaching an OD600 of 0.6–0.8 was reached. Protein production was induced by adding 20 µL of 1 M isopropyl β-D-1-thiogalactopyranoside (IPTG; Sigma), after which the culture was incubated overnight at 18 °C with shaking at 200 rpm. Cells were harvested the following day by centrifugation at 13,000 × g for 5 min at room temperature using an Eppendorf 5810R centrifuge. The pellet was resuspended in 50 mL of lysis buffer (50 mM Tris-HCl, 300 mM NaCl, 10 mM imidazole; pH 7.5), with 100 µL of 0.5 M phenylmethylsulfonyl fluoride (PMSF) and 100 µL of 1 M dithiothreitol (DTT) added. Cell lysis was performed by sonication using a Ultrasonic processor XL (Heat Systems) with following settings: maximum temperature 4 °C, amplitude 15 %, pulse on 10 s, pulse off 10 s, total lysis time of 10 min. The lysate was clarified by centrifugation at 13,000 rpm and 4 °C for 30 min in a Sorvall Lynx 6000 centrifuge (Thermo Scientific) using the F21 rotor. Samples were taken before and after centrifugation to check protein presence in the supernatant or pellet. For ion metal affinity purification, 1 mL of Ni-NTA agarose beads (Qiagen) were washed three times with 10 mL of lysis buffer. The supernatant was incubated with the washed beads for 1 h at 4 °C with rotation. The bead-protein mixture was transferred to a PD-10 poly-prep chromatography column (Bio-Rad) and washed with 15 mL of wash buffer (50 mM Tris-HCl, 300 mM NaCl, 20 mM imidazole; pH 7.5). The bound protein was eluted with 0.5 mL aliquots of elution buffer (50 mM Tris-HCl, 200 mM NaCl; pH 7.5) containing increasing concentrations of imidazole (50, 100, 150, and 200 mM). Eluted HisMBP-tagged protein samples (30 µL) were mixed with 10 µL 4x sample buffer (50 mM DTT in 4× Laemmli sample buffer; Bio-Rad), boiled at 95 °C for 5 minutes, and centrifuged at 13,000 rpm for 2 min. The mixture (20 µL) was loaded onto a Mini-Protein TGX Stain-Free Gel ((#4568083, Bio-Rad) and run at 160 V for 40 minu using a Mini-PROTEAN Tetra Vertical Electrophoresis Cell (Bio-Rad). The gel was stained with Instant Blue^TM^ for 4 h and then de-stained in water for another 4 h. After staining, gels were digitalized by scanning using an Expression 11000XL flatbed scanner (Epson). Protein concentrations were quantified using the Bradford assay. A standard calibration curve was generated from a serial dilution series of bovine serum albumin (BSA), and sample concentrations were calculated based on their absorbance values relative to this standard curve.

### Enzyme assays

#### PAL enzyme assays

Enzymatic assays employed 2 µg of purified enzyme and were conducted in 100 µL of 10 mM Tris–HCl buffer (pH 8.5) containing 1 mM L-phenylalanine and incubated at 37 °C for 4 h. All reactions were performed in replicates (n ≥ 3). Product formation was monitored continuously by measuring the absorbance at 270 nm every 5 min to quantify cinnamic acid formation using a SpectraMax ABS-Plus spectrophotometer (Molecular Devices).

#### TAL enzyme assays

Enzymatic assays employed 4 µg of purified enzyme and were conducted in 100 µL of 100 mM Tris–HCl buffer (pH 8) supplemented with 1 mM L-tyrosine, at 37 °C for 2 h. All reactions were performed in replicates (n ≥ 3). Product formation was monitored continuously by measuring the absorbance at 310 nm every 5 min to quantify *p*-coumaric acid formation using a SpectraMax ABS-Plus spectrophotometer (Molecular Devices).

#### CAL enzyme assays

Enzymatic assays employed 2 µg of purified enzyme and were conducted in 100 µL of 10 mM Tris-HCl buffer (pH 8.5), containing 1 mM 3-carboxy-tyrosine, at 37 °C for 3 h. All reactions were performed in replicates (n ≥ 3). The production of carboxy-*p*-coumaric acid was analysed by HPLC.

For the CAL enzyme kinetics, a concentration range between 0.146 μM and 1.25 mM 3-carboxy-tyrosine was used and the Michaelis–Menten equation was used to calculate K_M_, k_cat_ and V_max_ values. Kinetic data analysis was performed using GraphPad Prism10.

#### Activity on amino acid mixture

Purified PAL1 or PAL3 (1 µg) was used per reaction. The reactions were carried out in 100 µL of 100 mM Tris-HCl buffer (pH 8.5), containing 0.5 mM of each amino acid. Each reaction was performed in quadruplicate. The reactions were incubated for 30 min at 37 °C, whereupon 10 µL 6 M acetate was added to terminate the reaction. Samples were subjected to gas chromatography coupled to high resolution mass spectrometry (GC-HRMS) at the VIB Metabolomics Core Ghent. Briefly, samples were dried prior to derivatisation in a centrivap concentrator (Labconco) whereupon the metabolites were trimethylsilylated using 100 μL derivatisation mixture (N-methyl-N-(trimethylsilyl)trifluoroacetamide:pyridine in a 5:1 ratio). The samples were then shaken at 300 rpm for 1 h at room temperature. Prior to the analysis, the samples were diluted 1:50. GC-MS analyses were performed on an Agilent 7250 QTOF-MS, equipped with an Agilent 7890B GC system. One μL of derivatised sample was injected in splitless mode with the injector port set to 280 °C. Separation was achieved with a VF-5ms column (40 m x 0.25 mm; 0.25 μm; Varian CP9013; Agilent) with helium carrier gas at a constant flow rate of 1.2 mL/min. The oven was held at 80 °C for 1 min, ramped to 320 °C at a rate of 5 °C/min and held for 10 min. MS transfer line, MS ion source and quadrupole were held at 280 °C, 230 °C and 150 °C respectively. The MS detector was operated in electron ionization (EI) mode at 70 eV. Full EI-MS spectra were recorded by scanning the *m*/*z* range of 50 to 800 with a solvent delay of 7.8 min. For relative quantification, the peak areas were integrated using Agilent Masshunter Quantitative Analysis software. Identification of the amino acids was performed by comparison with the retention time and mass spectrum of reference standards.

### Metabolic profiling

Plants were grown on a vertical plate. At 14 DAS, roots were harvested and immediately frozen in liquid nitrogen. Five seedlings were pooled together per sample, and five and four biological replicates were used for WT and *pal* mutants (*pal3-3*, and *pal3-4)*, respectively. Subsequently, plant material was ground in 2-mL microcentrifuge tubes with precooled 4-mm stainless steel beads using a Retsch ball milling machine (20 Hz, 30 s). The phenolic metabolites were extracted with 1 mL methanol for 15 min at 60 °C in a thermomixer (1000 rpm). An 800-µL aliquot of each phenolic extract was dried at reduced pressure in a SpeedVac and subsequently solubilised in 100 µL cyclohexane and 100 µL MQ water. The water phase (15 µL) was injected on an ultra-high performance liquid chromatography (UHPLC) system (Waters Acquity UPLC^®^), equipped with a BEH C18 column (2.1 x 150 mm, 1.7 μM; Waters) and coupled to a quadrupole time-of-flight mass spectrometer [TOF MS, Synapt Q-Tof (Waters Corporation, Milford, Massachusetts, USA)]. The UHPLC buffer gradient and negative mode MS settings were as previously described (Hoengenaert et al. 2022). The ions eluting between 2 and 30 min were considered. Each peak intensity was normalized according to the total sum of the peak intensities in that sample. The following filters were applied for the untargeted analysis: (i) detected in all biological repeats of at least one genotype group, (ii) average ion intensity within at least one genotype group > 1000, (iii) ANOVA p-value < 0.0005, (iv) Student’s *t-*test p-value< 0.01, and (v) fold-change > 2 or < 0.5 in both lines compared to WT. Statistical analyses were performed on arcsinh transformed ion intensities. Structural elucidation was based on DDA-MSMS spectra where the low mass ramp was ramped between 15-35 eV and the high mass ramp was ramped between 35-70 eV with a scan time of 0.2 s.

### Molecular modelling and docking

Primary protein sequences of PAL3 obtained from TAIR (https://www.arabidopsis.org) were uploaded into AlphaFold Protein Structure Database (https://alphafold.ebi.ac.uk/). The tetrameric structures of PAL1 and PAL3 (apo-protein) were modelled using the AlphaFold3 server (Abramson et al. 2024). Additionally, an AlphaFill monomeric model was generated to determine an initial position for the MIO group (Hekkelman et al. 2023). Following this, the MIO forming residues (PAL1: A211, S212, and G213; PAL3: A191, S192, and G193) in chain A of the apo-protein were manually replaced with the MIO group obtained from the AlphaFill model (holo-monomer). For parametrization, the isolated MIO group was capped with an N-terminal acetate cap and a C-terminal N-methyl cap. AM1-BCC charges for this molecule were then calculated using “antechamber” (AmberTools suite version: 24.8) (Case et al. 2023). Hydrogens were added using “reduce” included in the AmberTools suite and correct protonation of titratable residues in the active site was verified using PROPKA at a pH of 7.4 (Søndergaard et al. 2011). The position of the MIO group in the holo-monomer was then dynamically tested with 1,000-steps of steepest descent and 1,000-steps of conjugate gradient minimization, using “sander” included in the AmberTools suite. During minimization all atoms, except for the MIO group, were constrained with a force of 10 kcal/mol. Finally, chain A of the previously generated tetrameric apo-protein was replaced with the minimized holo-monomer.

For molecular docking experiments, the MGLtools suite (version: 1.5.7) was used for the parametrization of ligands and receptors (Morris et al. 2009; Hekkelman et al. 2023). The python bindings of the Vina software (version 1.2.6) were used for docking, employing an exhaustiveness of 64 (Eberhardt et al. 2021). The cubic docking box was centered at the nucleophilic carbon of the MIO group with an equal edge length of 25 Å. Obtained, docking poses were visually analyzed using PyMOL (version: 3.1.0).

### Phylogenomic profiling of ammonia lyase-encoding genes

The PLAZA Dicots 5.0 and PLAZA Monocots 5.0 databases were searched (Van Bel et al. 2021) using the gene AT5G04230 (*AtPAL3*) as a query. The query gene was found in gene families HOM05D000844 and HOM05M000428 in PLAZA Dicots and PLAZA Monocots, respectively. To identify additional *PAL3* genes (i.e. containing amino acid residues corresponding to Y121, V448 and S472 in PAL3) within these gene families, a multiple sequence alignment (MSA) was performed of all these protein sequences via MAFFT (Katoh and Standley 2013) with parameters ‘--localpair --maxiterate 1000’. After trimming the MSA to remove columns with > 50 % of gaps, the positions of the catalytic pockets were identified by searching for motifs ‘LIRY’, ‘EVAM’, and ‘SASQ’ in the AT5G04230 sequence. PAL sequences with amino acid residues corresponding to PAL3 Y121 were classified as PAL3-like genes. PAL sequences with a histidine residue at the substrate specificity site were classified as PTALs.

To infer an ammonia lyase gene tree phylogeny, the trimmed MSA (see section above) was filtered to remove genes from species with no PAL3-like genes. The filtered MSA was used to infer an unrooted, maximum likelihood-based gene tree using IQ-TREE 2 (Minh et al. 2020) with 1000 bootstrap replicates. The tree was visualized using the ggtree R package (Yu et al. 2017).

## Notes

### Competing Interest Statement

The authors have declared no competing interest.

## REFERENCES

1 Parmeggiani, F., Weise, N. J., Ahmed, S. T. & Turner, N. J. Synthetic and Therapeutic Applications of Ammonia-lyases and Aminomutases. Chemical Reviews 118, 73–118 (2018). 10.1021/acs.chemrev.6b00824

2 Barros, J. & Dixon, R. A. Plant Phenylalanine/Tyrosine Ammonia-lyases. Trends in Plant Science 25, 66–79 (2020). 10.1016/j.tplants.2019.09.011

3 Verhoeven, J. et al. New fluoroethyl phenylalanine analogues as potential LAT1-targeting PET tracers for glioblastoma. Sci Rep 9, 2878 (2019). 10.1038/s41598-019-40013-x

4 Sun, C. et al. Direct asymmetric synthesis of β-branched aromatic α-amino acids using engineered phenylalanine ammonia lyases. Nature Communications 15, 8264 (2024). 10.1038/s41467-024-52613-x

5 Barros, J. et al. Role of bifunctional ammonia-lyase in grass cell wall biosynthesis. Nature Plants 2, 16050 (2016). 10.1038/nplants.2016.50

6 Boerjan, W., Ralph, J. & Baucher, M. Lignin biosynthesis. Annu. Rev. Plant Biol. 54, 519–546 (2003). 10.1146/annurev.arplant.54.031902.134938

7 Vogt, T. Phenylpropanoid biosynthesis. Molecular Plant 3, 2–20 (2010). 10.1093/mp/ssp106

8 Raes, J., Rohde, A., Christensen, J. H., Van de Peer, Y. & Boerjan, W. Genome-wide characterization of the lignification toolbox in Arabidopsis. Plant Physiol 133, 1051–1071 (2003). 10.1104/pp.103.026484

9 Rohde, A. et al. Molecular phenotyping of the *pal1* and *pal2* mutants of *Arabidopsis thaliana* reveals far-reaching consequences on phenylpropanoid, amino acid, and carbohydrate metabolism. Plant Cell 16, 2749–2771 (2004). 10.1105/tpc.104.023705

10 Huang, J. et al. Functional analysis of the Arabidopsis *PAL* gene family in plant growth, development, and response to environmental stress. Plant Physiology 153, 1526–1538 (2010). 10.1104/pp.110.157370

11 Leal, A. R. et al. Translational profile of developing phellem cells in Arabidopsis thaliana roots. The Plant Journal 110, 899–915 (2022).

12 Andersen, T. G. et al. Tissue-Autonomous Phenylpropanoid Production Is Essential for Establishment of Root Barriers. Curr Biol 31, 965–977.e965 (2021). 10.1016/j.cub.2020.11.070

13 Cochrane, F. C., Davin, L. B. & Lewis, N. G. The Arabidopsis phenylalanine ammonia lyase gene family: kinetic characterization of the four PAL isoforms. Phytochemistry 65, 1557–1564 (2004). 10.1016/j.phytochem.2004.05.006

14 Sun, L. et al. A dual role of Arabidopsis PAL nuclear localization in fine tuning flavonoid biosynthesis. Science Advances 12, eadz6970 (2026). doi:10.1126/sciadv.adz6970

15 Cooke, H. A., Christianson, C. V. & Bruner, S. D. Structure and chemistry of 4-methylideneimidazole-5-one containing enzymes. Curr Opin Chem Biol 13, 460–468 (2009). 10.1016/j.cbpa.2009.06.013

16 Watts, K. T., Mijts, B. N., Lee, P. C., Manning, A. J. & Schmidt-Dannert, C. Discovery of a Substrate Selectivity Switch in Tyrosine Ammonia-Lyase, a Member of the Aromatic Amino Acid Lyase Family. Chemistry & Biology 13, 1317–1326 (2006). 10.1016/j.chembiol.2006.10.008

17 Zoń, J. et al. Experimental and ab initio calculated structures of 2-aminoindane-2-phosphonic acid, a potent inhibitor of phenylalanine ammonia-lyase, and theoretical studies of its binding to the model enzyme structure. New Journal of Chemistry 28, 1048–1055 (2004).

18 Zoń, J., Miziak, P., Amrhein, N. & Gancarz, R. Inhibitors of phenylalanine ammonia-lyase (PAL): synthesis and biological evaluation of 5-substituted 2-aminoindane-2-phosphonic acids. Chem Biodivers 2, 1187–1194 (2005). 10.1002/cbdv.200590089

19 Zhao, T. et al. Whole-genome microsynteny-based phylogeny of angiosperms. Nature Communications 12, 3498 (2021). 10.1038/s41467-021-23665-0

20 Zhang, X. & Liu, C.-J. Multifaceted regulations of gateway enzyme phenylalanine ammonia-lyase in the biosynthesis of phenylpropanoids. Molecular Plant 8, 17–27 (2015). 10.1016/j.molp.2014.11.001

21 Larsen, P. O. m-Carboxy-substituted aromatic amino acids in plant metabolism: II. The incorporation of shikimic acid into L-3-(3-carboxy-4-hydroxyphenyl)-alanine in Reseda lutea L. Biochimica et Biophysica Acta (BBA) -General Subjects 115, 529–531 (1966). 10.1016/0304-4165(66)90463-6

22 Larsen, P. O. Meta-carboxy-substituted aromatic amino acids in plant metabolism. 3. Biochim Biophys Acta 141, 27–46 (1967). 10.1016/0304-4165(67)90242-5

23 Wu, J. et al. The cytosolic aminotransferase VAS1 coordinates aromatic amino acid biosynthesis and metabolism. Science Advances 10, eadk0738 (2024).

24 Wildermuth, M. C., Dewdney, J., Wu, G. & Ausubel, F. M. Isochorismate synthase is required to synthesize salicylic acid for plant defence. Nature 414, 562–565 (2001). 10.1038/35107108

25 Garcion, C. et al. Characterization and biological function of the ISOCHORISMATE SYNTHASE2 gene of Arabidopsis. Plant Physiol 147, 1279–1287 (2008). 10.1104/pp.108.119420

26 Thilmony, R., Underwood, W. & He, S. Y. Genome-wide transcriptional analysis of the Arabidopsis thaliana interaction with the plant pathogen Pseudomonas syringae pv. tomato DC3000 and the human pathogen Escherichia coli O157: H7. The Plant Journal 46, 34–53 (2006).

27 Guo, W. et al. An ethylene response-related factor, GbERF1-like, from Gossypium barbadense improves resistance to Verticillium dahliae via activating lignin synthesis. Plant Mol Biol 91, 305–318 (2016). 10.1007/s11103-016-0467-6

## References

Abramson J, Adler J, Dunger J, Evans R, Green T, Pritzel A, Ronneberger O, Willmore L, Ballard AJ, Bambrick J, Bodenstein SW, Evans DA, Hung C-C, O’Neill M, Reiman D, Tunyasuvunakool K, Wu Z, Žemgulytė A, Arvaniti E, Beattie C, Bertolli O, Bridgland A, Cherepanov A, Congreve M, Cowen-Rivers AI, Cowie A, Figurnov M, Fuchs FB, Gladman H, Jain R, Khan YA, Low CMR, Perlin K, Potapenko A, Savy P, Singh S, Stecula A, Thillaisundaram A, Tong C, Yakneen S, Zhong ED, Zielinski M, Žídek A, Bapst V, Kohli P, Jaderberg M, Hassabis D, Jumper JM (2024) Accurate structure prediction of biomolecular interactions with AlphaFold 3. Nature 630 (8016):493–500. doi:10.1038/s41586-024-07487-w

Arnold ZaL, P. O. (1977) A New Synthesis of 3-(3-Carboxy-4-hydroxyphenyl)-L-alanine (3′-Carboxy-L-tyrosine). Acta Chemica Scandinavica, Series B 31:826–828. doi:10.3891/acta.chem.scand.31b-0826

Arnold ZS (1985) Optically active aromatic amino acids. Part IV. Synthesis and analgesic activity of four derivatives of D-Tyrosine. Polish Journal of Chemistry 59:837–843

Case DA, Aktulga HM, Belfon K, Cerutti DS, Cisneros GA, Cruzeiro VWD, Forouzesh N, Giese TJ, Götz AW, Gohlke H, Izadi S, Kasavajhala K, Kaymak MC, King E, Kurtzman T, Lee T-S, Li P, Liu J, Luchko T, Luo R, Manathunga M, Machado MR, Nguyen HM, O’Hearn KA, Onufriev AV, Pan F, Pantano S, Qi R, Rahnamoun A, Risheh A, Schott-Verdugo S, Shajan A, Swails J, Wang J, Wei H, Wu X, Wu Y, Zhang S, Zhao S, Zhu Q, Cheatham TE, III, Roe DR, Roitberg A, Simmerling C, York DM, Nagan MC, Merz KM, Jr. (2023) AmberTools. Journal of Chemical Information and Modeling 63 (20):6183–6191. doi:10.1021/acs.jcim.3c01153

Eberhardt J, Santos-Martins D, Tillack AF, Forli S (2021) AutoDock Vina 1.2.0: New Docking Methods, Expanded Force Field, and Python Bindings. Journal of Chemical Information and Modeling 61 (8):3891–3898. doi:10.1021/acs.jcim.1c00203

Hekkelman ML, de Vries I, Joosten RP, Perrakis A (2023) AlphaFill: enriching AlphaFold models with ligands and cofactors. Nature Methods 20 (2):205–213. doi:10.1038/s41592-022-01685-y

Hoengenaert L, Wouters M, Kim H, De Meester B, Morreel K, Vandersyppe S, Pollier J, Desmet S, Goeminne G, Ralph J, Boerjan W, Vanholme R (2022) Overexpression of the scopoletin biosynthetic pathway enhances lignocellulosic biomass processing. Sci Adv 8 (28):eabo5738. doi:10.1126/sciadv.abo5738

Leroux M, Vorherr T, Lewis I, Schaefer M, Koch G, Karaghiosoff K, Knochel P (2019) Late-Stage Functionalization of Peptides and Cyclopeptides Using Organozinc Reagents. Angew Chem Int Ed Engl 58 (24):8231–8234. doi:10.1002/anie.201902454

Lokolkar M, Bhanage B (2023) Palladium Catalysed Carbonylative Homocoupling of 2-iodophenols for Synthesis of Symmetrical Xanthones. Synlett 0. doi:10.1055/a-2123-9288

Morris GM, Huey R, Lindstrom W, Sanner MF, Belew RK, Goodsell DS, Olson AJ (2009) AutoDock4 and AutoDockTools4: Automated docking with selective receptor flexibility. J Comput Chem 30 (16):2785–2791. doi:10.1002/jcc.21256

Schneider CA, Rasband WS, Eliceiri KW (2012) NIH Image to ImageJ: 25 years of image analysis. Nat Methods 9 (7):671–675. doi:10.1038/nmeth.2089

Søndergaard CR, Olsson MH, Rostkowski M, Jensen JH (2011) Improved Treatment of Ligands and Coupling Effects in Empirical Calculation and Rationalization of pKa Values. J Chem Theory Comput 7 (7):2284–2295. doi:10.1021/ct200133y

