## Supplementary material for "The Arabidopsis PAL3 is a carboxy-tyrosine ammonia lyase": Suppl Figures

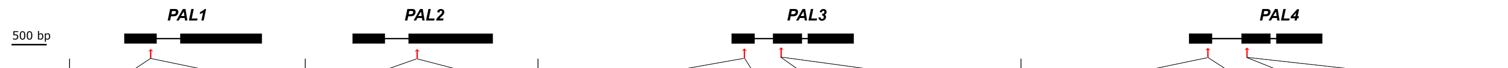


| | PAL1 | PAL2 | PAL3 | | PAL4 | |
|---|---|---|---|---|---|---|
| WT | GATCGCAGCCACTTGTCCAATGG | GTTGACTCTGACGAGCATGGCGG | GTTGCGGCAGTAGCGAGTGGAGG | GTTCCGTTATCCTATATCGCTGG | GTGATGGAGAGCATGAACCGAGG | GTGACGTGTCCCCGGCGCCGGG |
| *pal1-4* | +A: GATCGCAGCCACTTGTCACAATGG | | | | | |
| *pal1-5* | +C: GATCGCAGCCACTTGTCCCAATGG | | | | | |
| *pal2-4* | | -A: GTTGACTCTGACGAGC-TGGCGG | | | | |
| *pal2-5* | | +A: GTTGACTCTGACGAGCAATGGCGG | | | | |
| *pal3-3* | | | -G: GTTGCGGCAGTAGCGA-TGGAGG | +A: GTTCCGTTATCCTATATACGCTGG | | |
| *pal3-4* | | | +G: GTTGCGGCAGTAGCGAGGTGGAGG | 0: GTTCCGTTATCCTATATCGCTGG | | |
| *pal4-3* | | | | | +A: GTGATGGAGAGCATGAAACCGAGG | +A: GTGACGTGTCCCCGGCGACCGGG |
| *pal4-4* |  |  |  |  | -A: GTGATGGAGAGCATGA-CCGAGG | -38: (-3)--------------------(-12) |
| *pal1/2-4* | +A: GATCGCAGCCACTTGTCACAATGG | -A: GTTGACTCTGACGAGC-TGGCGG | | | | |
| *pal1/2-5* | +C: GATCGCAGCCACTTGTCCCAATGG | +A: GTTGACTCTGACGAGCAATGGCGG | | | | |
| *pal1/2/3-3* | +A: GATCGCAGCCACTTGTCACAATGG | -A: GTTGACTCTGACGAGC-TGGCGG | +G: GTTGCGGCAGTAGCGAGGTGGAGG | +A: GTTCCGTTATCCTATATACGCTGG | | |
| *pal1/2/3-5* | +C: GATCGCAGCCACTTGTCCCAATGG | +A: GTTGACTCTGACGAGCAATGGCGG | -G: GTTGCGGCAGTAGCGA-TGGAGG | 0: GTTCCGTTATCCTATATCGCTGG | | |
| *pal1/2/4-3* | +A: GATCGCAGCCACTTGTCACAATGG | -A: GTTGACTCTGACGAGC-TGGCGG | | | +A: GTGATGGAGAGCATGAAACCGAGG | +A: GTGACGTGTCCCCGGCGACCGGG |
| *pal1/2/4-5* | +C: GATCGCAGCCACTTGTCCCAATGG | +A: GTTGACTCTGACGAGCAATGGCGG |  |  | -A: GTGATGGAGAGCATGA-CCGAGG | -38: (-3)--------------------(-12) |
| *pal1/2/3/4-3* | +C: GATCGCAGCCACTTGTCCCAATGG | +A: GTTGACTCTGACGAGCAATGGCGG | +G: GTTGCGGCAGTAGCGAGGTGGAGG | +A: GTTCCGTTATCCTATATACGCTGG | -A: GTGATGGAGAGCATGA-CCGAGG | -38: (-3)--------------------(-12) |

Suppl Fig 1

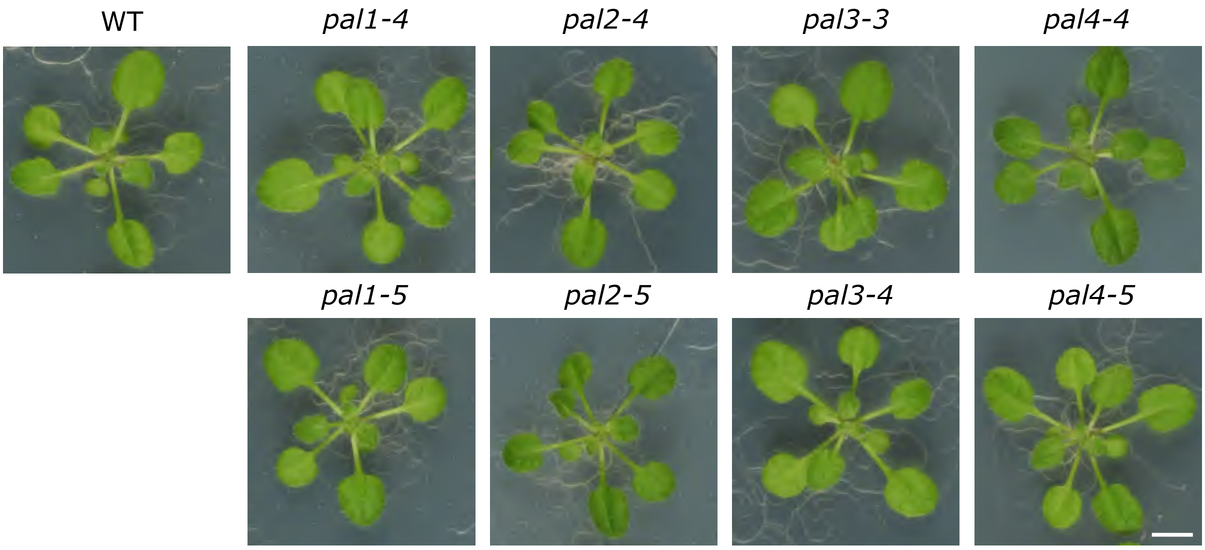


Suppl Fig 2

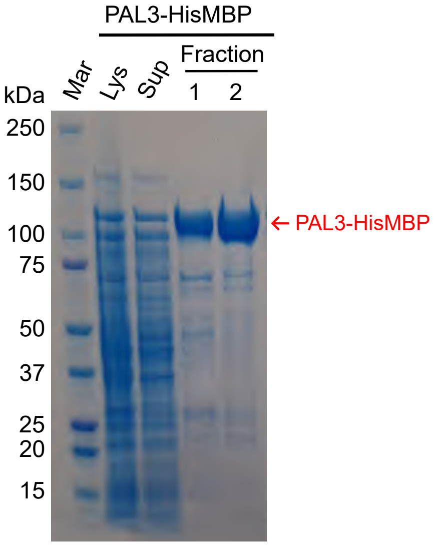
PAL3-HisMBP
Fraction
kDa
Mar
Lys
Sup
1
2
250
150
100
75
50
37
25
20
15
← PAL3-HisMBP

Supp Fig 3

```
                                                                                                                                                     L118
                                                                                                                                                        Y121
                                                                                                                          Y94                              L122
AtPAL1     MEINGAHKSNGGGVDAMLCGGDIKTKNMVINA--EDPLNWGAAAEQMKGSHLDEVKRMVAEFRKPVVNLGGETLTIGQVAAISTIGNSVKVELSETARAGVNASSDWVMESMNKGTDSYGVTTGFGATSHRRTKNGVALQKELIRFLNAG
AtPAL2     ----------MDQIEAMLCGGGEKTKVAVTTKTLADPLNWGLAADQMKGSHLDEVKKMVEEYRRPVVNLGGETLTIGQVAAISTVGGSVKVELAETSRAGVKASSDWVMESMNKGTDSYGVTTGFGATSHRRTKNGTALQTELIRFLNAG
AtPAL3     ----------------------MEFRQPNATA-LSDPLNWNVAAEALKGSHLEEVKKMVKDYRKGTVQLGGETLTIGQVAAVAS--GGPTVELSEEARGGVKASSDWVMESMNRDTDTYGITTGFGSSSRRRTDQGAALQKELIRYLNAG
AtPAL4     --------------------MELCNQNNHITAVSGDPLNWNATAEALKGSHLDEVKRMVKEYRKEAVKLGGETLTIGQVAAVARGGGGSTVELAEEARAGVKASSEWVMESMNRGTDSYGVTTGFGATSHRRTKQGGALQNELIRFLNAG

                                                                                                                           L244
                                                                                                                              V247
                                                                   MIO L195                                                    N248
AtPAL1     IF--GSTKE-TSHTLPHSATRAAMLVRINTLLQGFSGIRFEILEAITSFLNNNITPSLPLRGTITASGDLVPLSYIAGLLTGRPNSKATGPNGEALTAEEAFKLAGISSGFFDLQPKEGLALVNGTAVGSGMASMVLFETNVLSVLAEIL
AtPAL2     IF--GNTKE-TCHTLPQSATRAAMLVRVNTLLQGYSGIRFEILEAITSLLNHNISPSLPLRGTITASGDLVPLSYIAGLLTGRPNSKATGPDGESLTAKEAFEKAGISTGFFDLQPKEGLALVNGTAVGSGMASMVLFEANVQAVLAEVL
AtPAL3     IFATGNEDDDRSNTLPRPATRAAMLIRVNTLLQGYSGIRFEILEAITTLLNCKITPLLPLRGTITASGDLVPLSYIAGFLIGRPNSRSVGPSGEILTALEAFKLAGVSS-FFELRPKEGLALVNGTAVGSALASTVLYDANILVVFSEVA
AtPAL4     IF--GPGAGDTSHTLPKPTTRAAMLVRVNTLLQGYSGIRFEILEAITKLLNHEITPCLPLRGTITASGDLVPLSYIAGLLTGRPNSKAVGPSGETLTASEAFKLAGVSS-FFELQPKEGLALVNGTAVGSGLASTVLFDANILAVLSEVM

                                                                   Y339
                                                                      R342                                          F338
AtPAL1     SAVFAEVMSGKPEFTDHLTHRLKHHPGQIEAAAIMEHILDGSSYMKLAQKLHEMDPLQKPKQDRYALRTSPQWLGPQIEVIRYATKSIEREINSVNDNPLIDVSRNKAIHGGNFQGTPIGVSMDNTRLAIAAIGKLMFAQFSELVNDFYN
AtPAL2     SAIFAEVMSGKPEFTDHLTHRLKHHPGQIEAAAIMEHILDGSSYMKLAQKVHEMDPLQKPKQDRYALRTSPQWLGPQIEVIRQATKSIEREINSVNDNPLIDVSRNKAIHGGNFQGTPIGVSMDNTRLAIAAIGKLMFAQFSELVNDFYN
AtPAL3     SAMFAEVMQGKPEFTDHLTHKLKHHPGQIEAAAIMEHILDGSSYVKEALHLHKIDPLQKPKQDRYALRTSPQWLGPQIEVIRAATKMIEREINSVNDNPLIDVSRNKAIHGGNFQGTPIGVAMDNTRLALASIGKLMFAQFTELVNDFYN
AtPAL4     SAMFAEVMQGKPEFTDHLTHKLKHHPGQIEAAAIMEHILDGSSYVKEAQLLHEMDPLQKPKQDRYALRTSPQWLGPQIEVIRAATKMIEREINSVNDNPLIDVSRNKALHGGNFQGTPIGVAMDNSRLAIASIGKLMFAQFSELVNDFYN

                          K444
                             V448                S472
AtPAL1     NGLPSNLTASRNPSLDYGFKGAEIAMASYCSELQYLANPVTSHVQSAEQHNQDVNSLGLISSRKTSEAVDILKLMSTTFLVAICQAVDLRHLEENLRQTVKNTVSQVAKKVLTTGVNGELHPSRFCEKDLLKVVDREQVYTYADDPCSAT
AtPAL2     NGLPSNLTASSNPSLDYGFKGAEIAMASYCSELQYLANPVTSHVQSAEQHNQDVNSLGLISSRKTSEAVDILKLMSTTFLVGICQAVDLRHLEENLRQTVKNTVSQVAKKVLTTGINGELHPSRFCEKDLLKVVDREQVFTYVDDPCSAT
AtPAL3     NGLPSNLSGGRNPSLDYGLKGAEVAMASYCSELQFLANPVTNHVESASQHNQDVNSLGLISSRTTAEAVVILKLMSTTYLVALCQAFDLRHLEEILKKAVNEVVSHTAKSVLA------IEPFR-KHDDILGVVNREYVFSYVDDPSSLT
AtPAL4     NGLPSNLSGGRNPSLDYGFKGAEIAMASYCSELQFLANPVTNHVQSAEQHNQDVNSLGLISSRKTAEAVDILKLMSTTYLVALCQAVDLRHLEENLKKAVKSAVSQVAKRVLTVGANGELHPSRFTERDVLQVVDREYVFSYADDPCSLT

AtPAL1     YPLIQKLRQVIVDHALINGESEKNAVTSIFHKIGAFEEELKAVLPKEVEAARAAYDNGTSAIPNRIKECRSYPLYRFVREELGTELLTGEKVTSPGEEFDKVFTAICEGKIIDPMMECLNEWNGAPIPIC
AtPAL2     YPLMQRLRQVIVDHALSNGETEKNAVTSIFQKIGAFEEELKAVLPKEVEAARAAYGNGTAPIPNRIKECRSYPLYRFVREELGTKLLTGEKVVSPGEEFDKVFTAMCEGKLIDPLMDCLKEWNGAPIPIC
AtPAL3     NPLMQKLRHVLFDKALAEPEGETD---TVFRKIGAFEAELKFLLPKEVERVRTEYENGTFNVANRIKKCRSYPLYRFVRNELETRLLTGEDVRSPGEDFDKVFRAISQGKLIDPLFECLKEWNGAPISIC
AtPAL4     YPLMQKLRHILVDHALADPEREANSATSVFHKIGAFEAELKLLLPKEVERVRVEYEEGTSAIANRIKECRSYPLYRFVRDELNTELLTGENVRSPGEEFDKVFLAISDGKLIDPLLECLKEWNGAPVSIC
```

Supp Fig 4

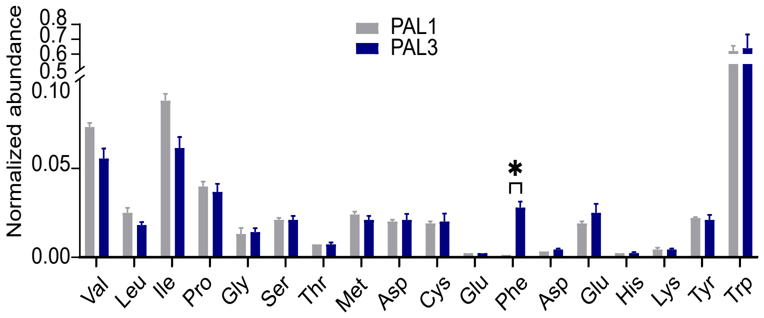

Normalized abundance
0.8
0.7
0.6
0.5
0.10
0.05
0.00
PAL1
PAL3
*
Val
Leu
Ile
Pro
Gly
Ser
Thr
Met
Asp
Cys
Glu
Phe
Asp
Glu
His
Lys
Tyr
Trp


Supp Fig 5

**A** AtGenExpress eFP browser - AT5G04230

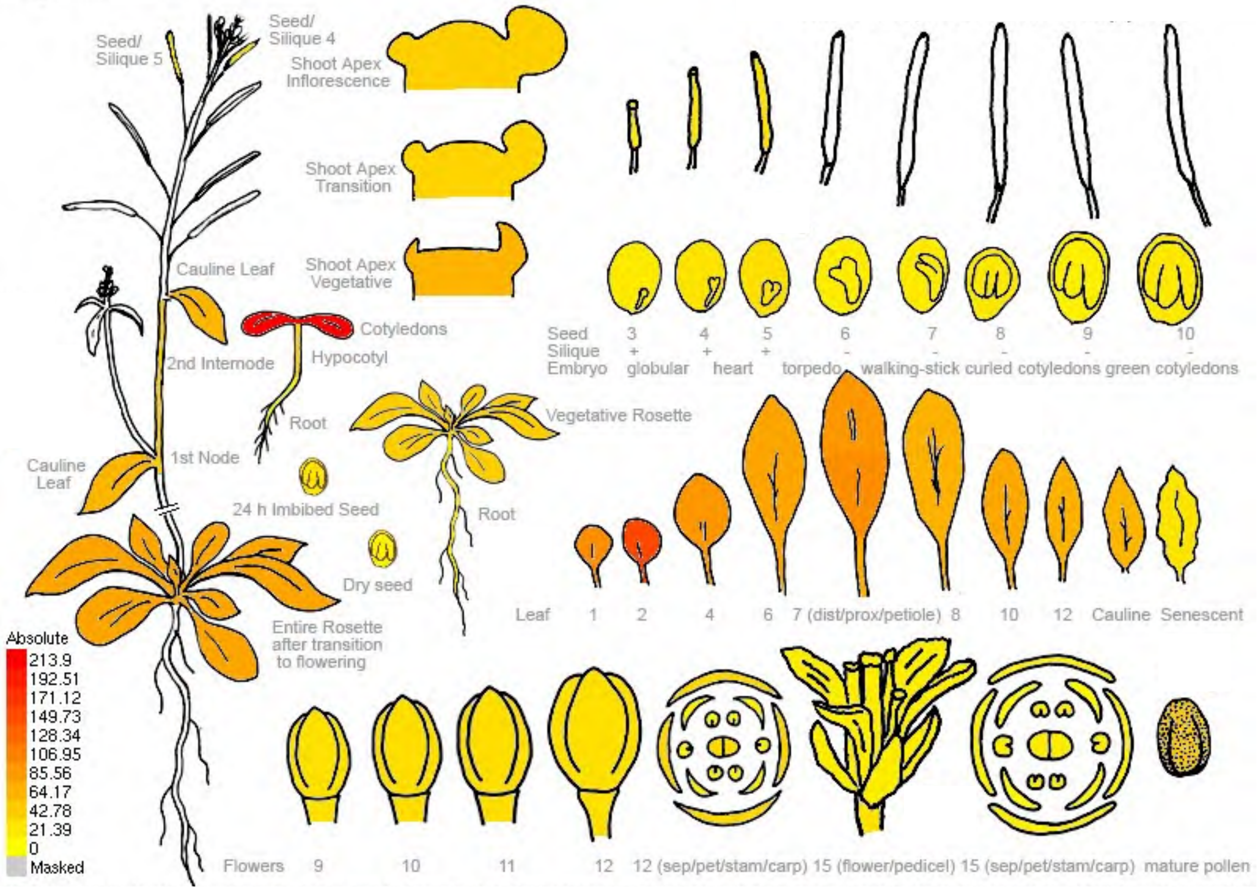


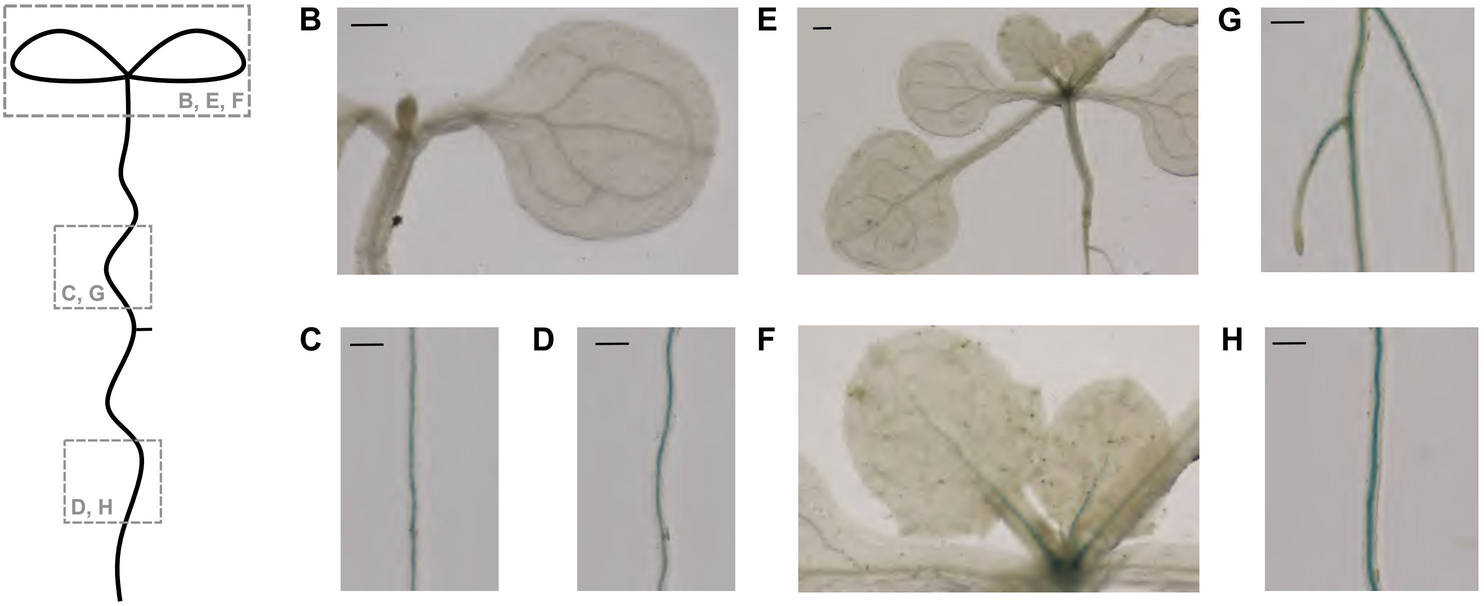


Supp Fig 6

### **Supplemental figure S7.** Structural elucidation of metabolites via MS/MS spectra

#### **2) 3-carboxy-*p*-coumaric acid + hexose 1**, m/z 369.0822, 10.14 min

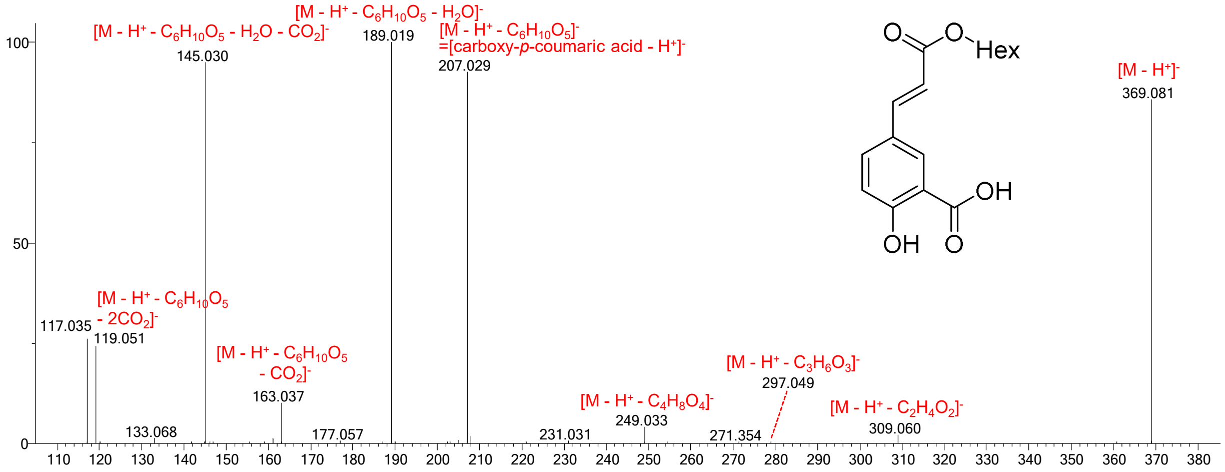


The $C_6H_{10}O_5$ neutral loss is dehydrated hexoside. Neutral losses of $C_2H_4O_2$, $C_3H_6O_3$, $C_4H_8O_4$ are part the hexose moiety. These losses, together with the combined loss of $C_6H_{10}O_6$ and $H_2O$, indicate that the sugar moiety is linked to one of the carboxyl groups, and not on the phenolic alcohol.

#### **3) 3-carboxy-*p*-coumaric acid + hexose 2**, m/z 369.0812, 10.54 min

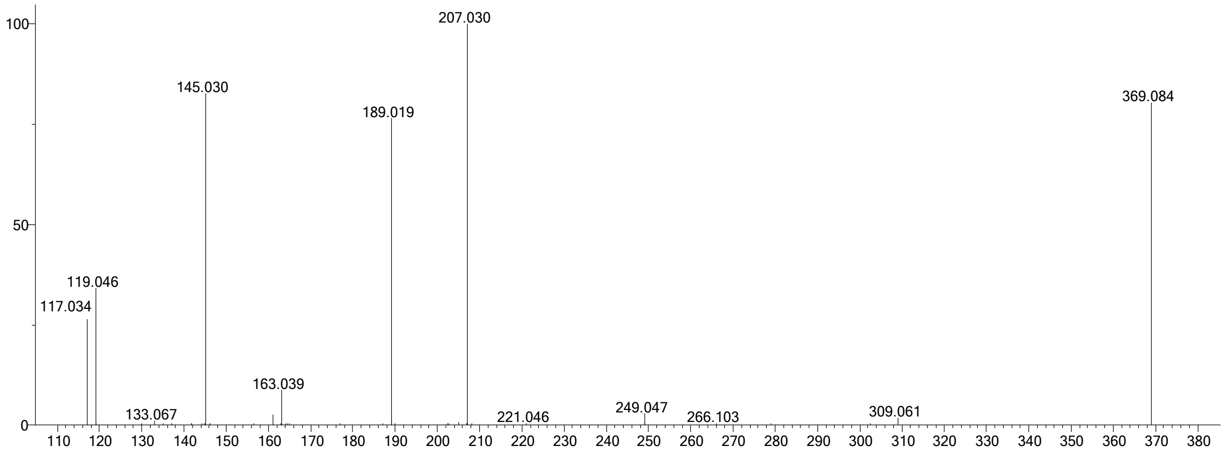


For structural elucidation, see compound **2**, 3-carboxy-*p*-coumaric acid + hexose 1.

#### **4) 3-carboxy-*p*-coumaric acid + lysine 1,** m/z 335.1242, 8.21 min no MS/MS available
#### **5) 3-carboxy-*p*-coumaric acid + lysine 2,** m/z 335.1281, 8.97 min

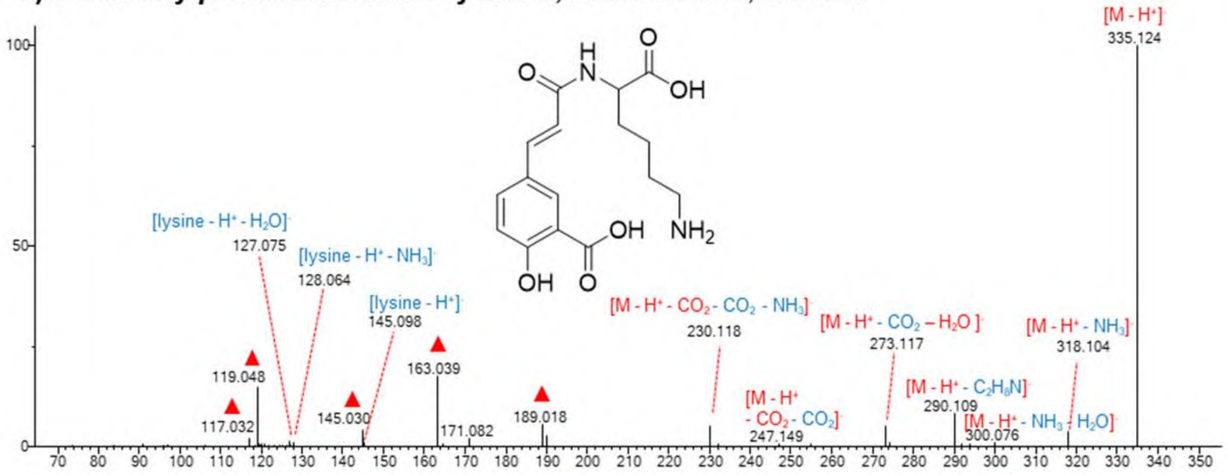


The ions and neutral losses derived from lysine are indicated in blue. Fragments indicative for 3-carboxy-p-coumaric acid (see compound **1** and **2**) are indicated with a red triangle▲.

### 6) **3-carboxy-*p*-coumaric acid + putrescine**, m/z 277.1206, 8.17 min

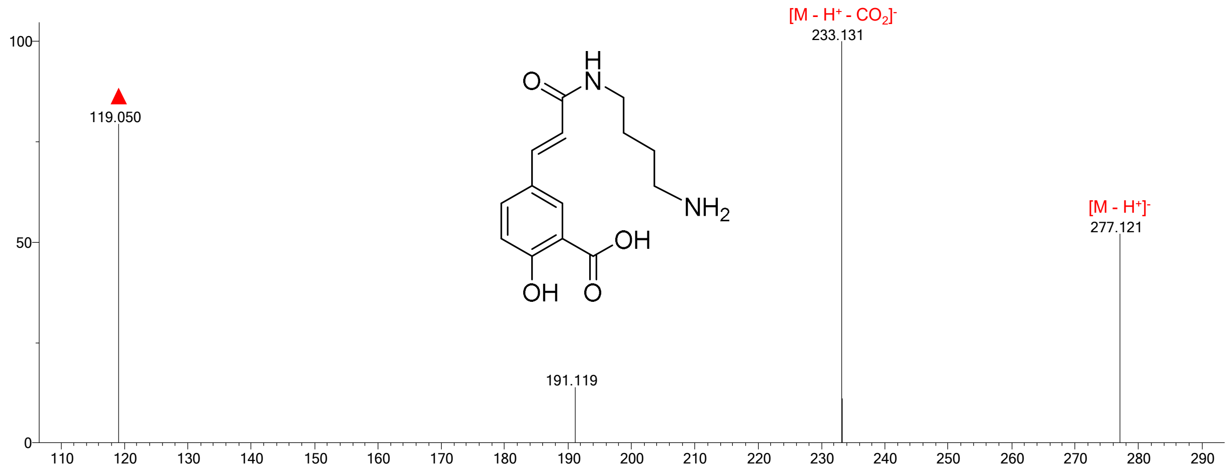


Note: the MS/MS spectrum is not of the highest quality. Nevertheless, the $CO_2$ neutral loss and a m/z 119, indicated with a red triangle are indicative for 3-carboxy-*p*-coumaric acid (see compound **1** and **2**) and clearly detected. The origin of m/z 191.119, matches $C_{11}H_{15}N_2O^-$ and is likely the result of a rearrangement.

### 7) **3-carboxycaffeic + hexose 1**, m/z 385.0764, 5.85 min

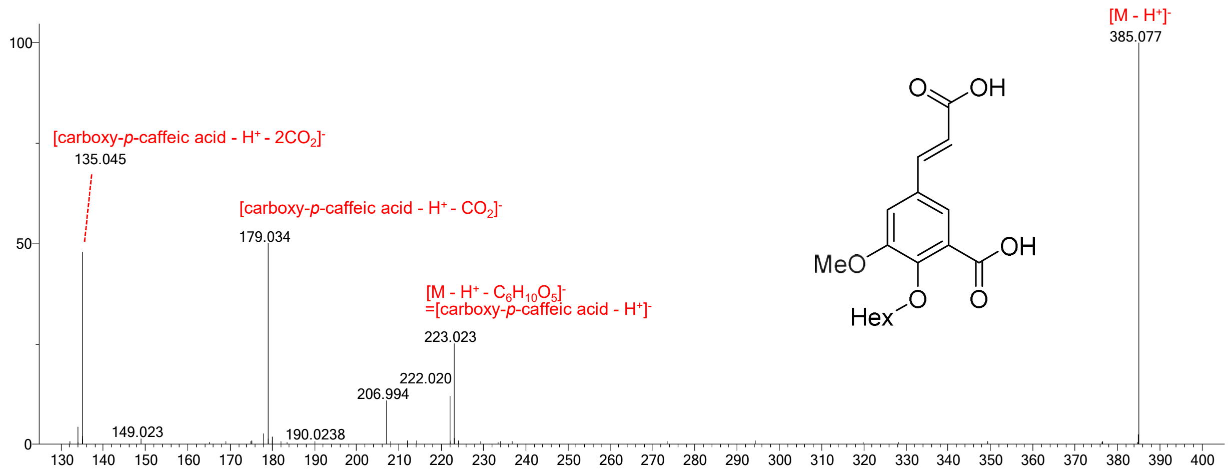


The $C_6H_{10}O_5$ neutral loss is dehydrated hexoside. The absence of neutral losses of $C_2H_4O_2$, $C_3H_6O_3$, $C_4H_8O_4$ (that are part the hexose moiety) and the absence of the combined loss of $C_6H_{10}O_6$ and $H_2O$, hint that the sugar moiety is linked to one of the phenolic groups, and not on the carboxylic groups.

### 8) **3-carboxycaffeic acid + hexose 2,** m/z 385.0766, 7.33 min

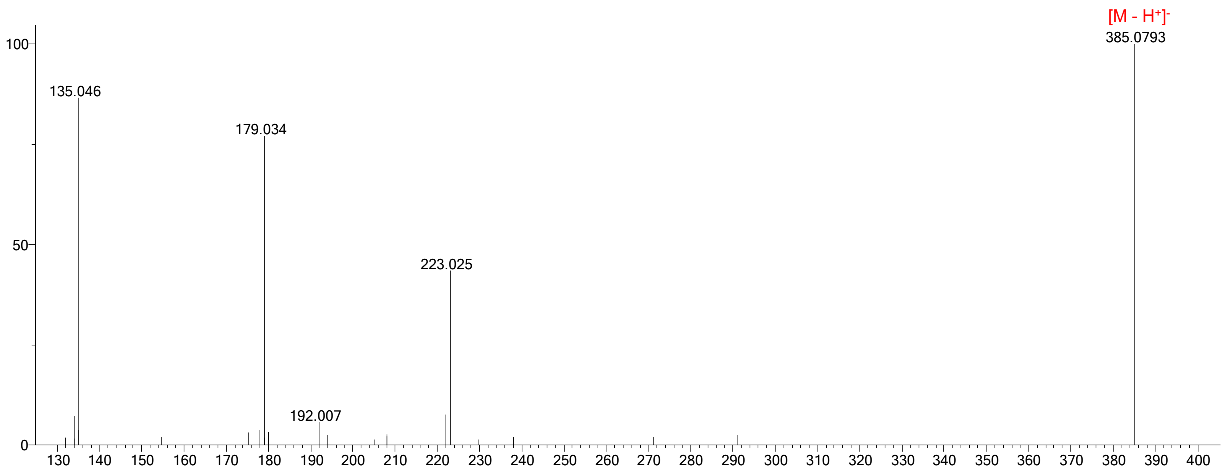


For structural elucidation, see compound **7**, 3-carboxycaffeic acid + hexose 1.

Supp Fig 7

**9) 3-carboxyferulic acid 1**, m/z 237.0386, 12.07 min

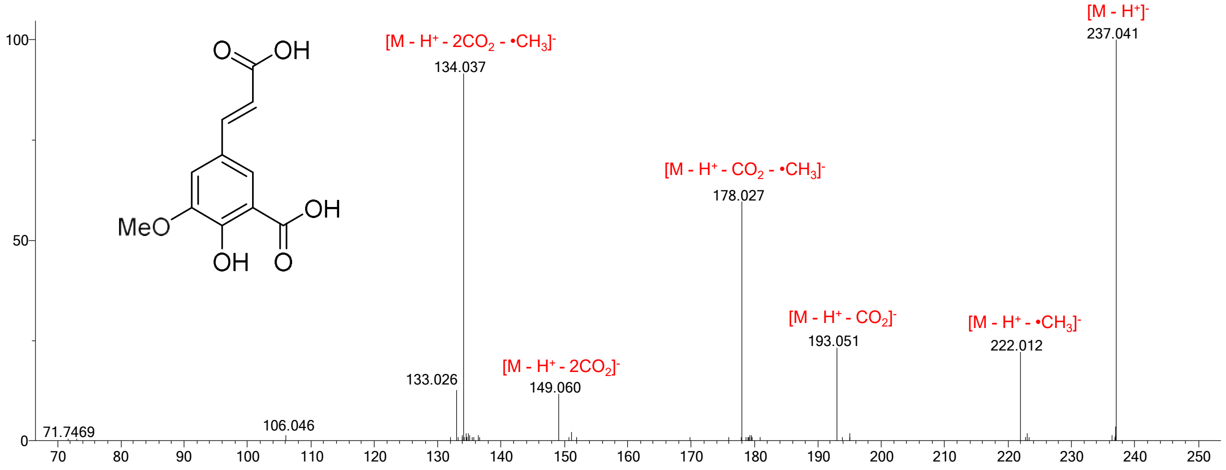


**10) 3-carboxyferulic acid 2**, m/z 237.0387, 12.73 min no MS/MS available

**3-carboxytyrosine**, m/z 224.0552, 4.38 min

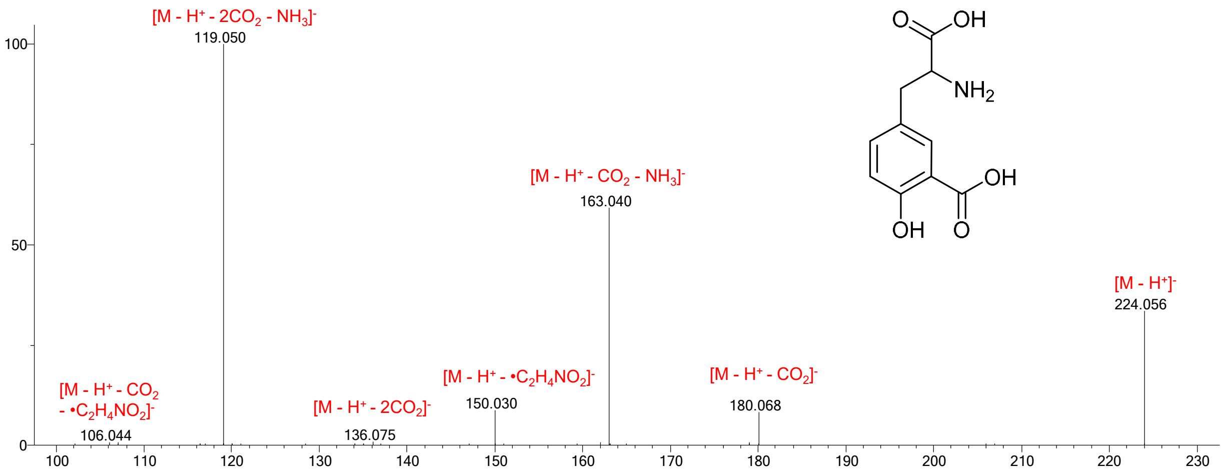


**3-carboxyphenylalanine**, m/z 208.0612, 3.76 min

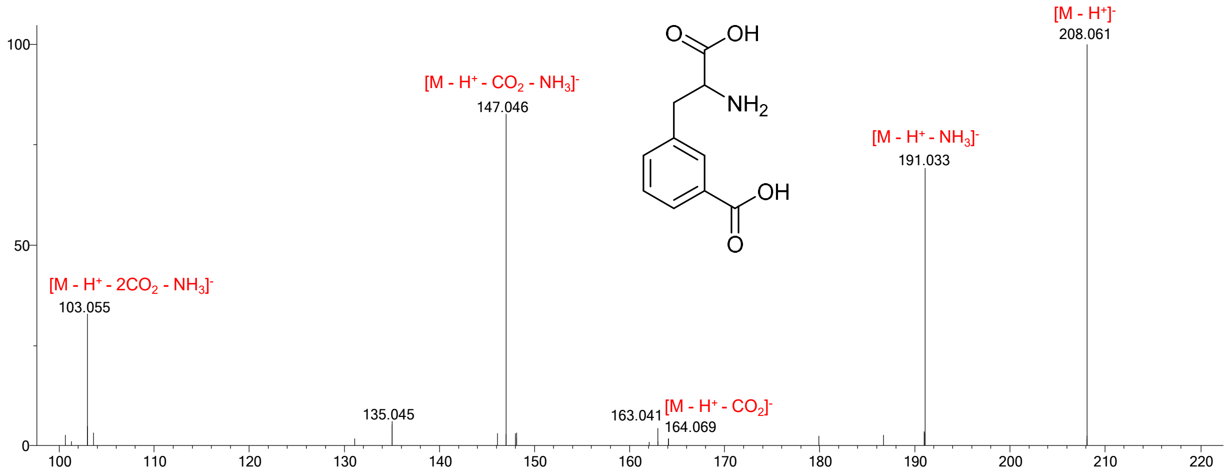


Supp Fig 7

A

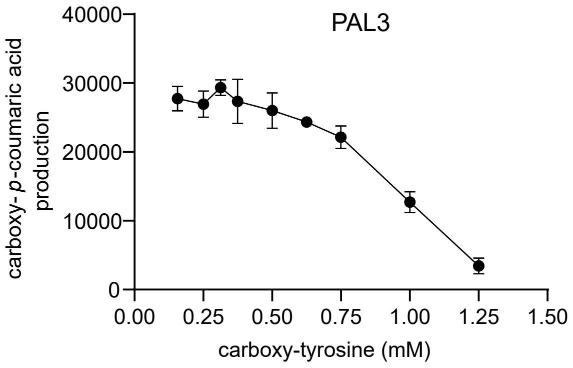
PAL3
40000
30000
20000
10000
0
carboxy-p-coumaric acid production
0.00
0.25
0.50
0.75
1.00
1.25
1.50
carboxy-tyrosine (mM)

B

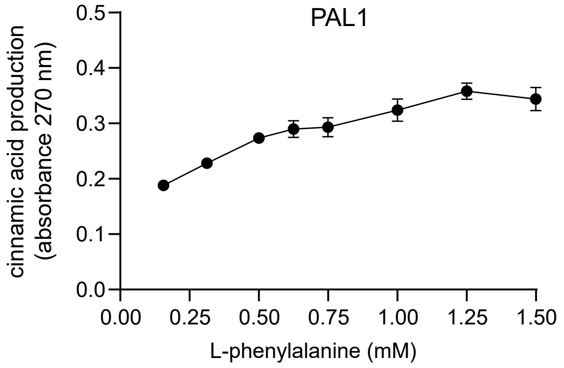
PAL1
0.5
0.4
0.3
0.2
0.1
0.0
cinnamic acid production (absorbance 270 nm)
0.00
0.25
0.50
0.75
1.00
1.25
1.50
L-phenylalanine (mM)

Supp Fig 8

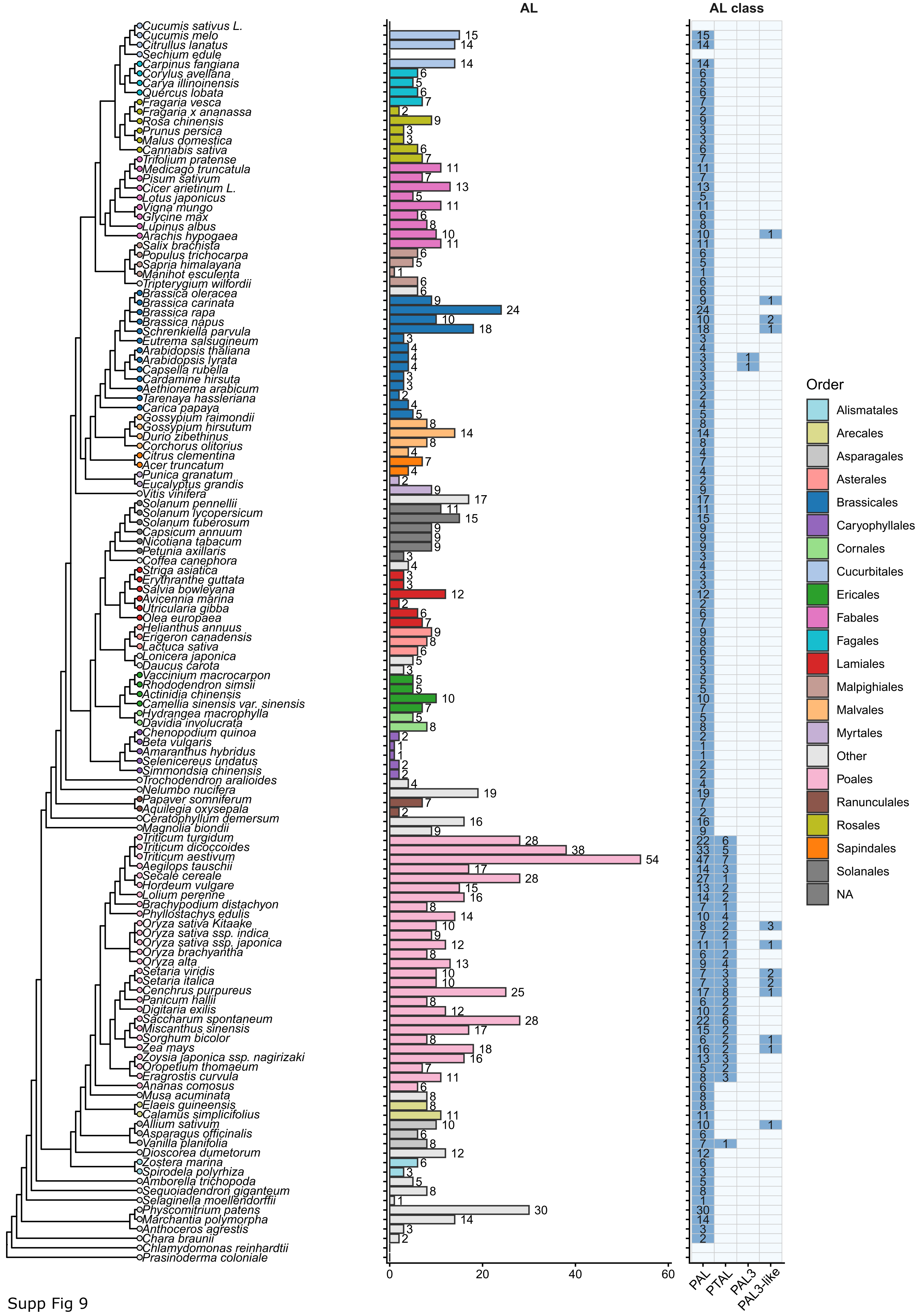
AL
AL class
PAL
PTAL
PAL3
PAL3-like
Order
Alismatales
Arecales
Asparagales
Asterales
Brassicales
Caryophyllales
Cornales
Cucurbitales
Ericales
Fabales
Fagales
Lamiales
Malpighiales
Malvales
Myrtales
Other
Poales
Ranunculales
Rosales
Sapindales
Solanales
NA
0
20
40
60
Cucumis sativus L.
Cucumis melo
Citrullus lanatus
Sechium edule
Carpinus fangiana
Corylus avellana
Carya illinoinensis
Quercus lobata
Fragaria vesca
Fragaria x ananassa
Rosa chinensis
Prunus persica
Malus domestica
Cannabis sativa
Trifolium pratense
Medicago truncatula
Pisum sativum
Cicer arietinum L.
Lotus japonicus
Vigna mungo
Glycine max
Lupinus albus
Arachis hypogaea
Salix brachista
Populus trichocarpa
Sapria himalayana
Manihot esculenta
Tripterygium wilfordii
Brassica oleracea
Brassica carinata
Brassica rapa
Brassica napus
Schrenkiella parvula
Eutrema salsugineum
Arabidopsis thaliana
Arabidopsis lyrata
Capsella rubella
Cardamine hirsuta
Aethionema arabicum
Tarenaya hassleriana
Carica papaya
Gossypium raimondii
Gossypium hirsutum
Durio zibethinus
Corchorus olitorius
Citrus clementina
Acer truncatum
Punica granatum
Eucalyptus grandis
Vitis vinifera
Solanum pennellii
Solanum lycopersicum
Solanum tuberosum
Capsicum annuum
Nicotiana tabacum
Petunia axillaris
Coffea canephora
Striga asiatica
Erythranthe guttata
Salvia bowleyana
Avicennia marina
Utricularia gibba
Olea europaea
Helianthus annuus
Erigeron canadensis
Lactuca sativa
Lonicera japonica
Daucus carota
Vaccinium macrocarpon
Rhododendron simsii
Actinidia chinensis
Camellia sinensis var. sinensis
Hydrangea macrophylla
Davidia involucrata
Chenopodium quinoa
Beta vulgaris
Amaranthus hybridus
Selenicereus undatus
Simmondsia chinensis
Trochodendron aralioides
Nelumbo nucifera
Papaver somniferum
Aquilegia oxysepala
Ceratophyllum demersum
Magnolia biondii
Triticum turgidum
Triticum dicoccoides
Triticum aestivum
Aegilops tauschii
Secale cereale
Hordeum vulgare
Lolium perenne
Brachypodium distachyon
Phyllostachys edulis
Oryza sativa Kitaake
Oryza sativa ssp. indica
Oryza sativa ssp. japonica
Oryza brachyantha
Oryza alta
Setaria viridis
Setaria italica
Cenchrus purpureus
Panicum hallii
Digitaria exilis
Saccharum spontaneum
Miscanthus sinensis
Sorghum bicolor
Zea mays
Zoysia japonica ssp. nagirizaki
Oropetium thomaeum
Eragrostis curvula
Ananas comosus
Musa acuminata
Elaeis guineensis
Calamus simplicifolius
Allium sativum
Asparagus officinalis
Vanilla planifolia
Dioscorea dumetorum
Zostera marina
Spirodela polyrhiza
Amborella trichopoda
Sequoiadendron giganteum
Selaginella moellendorffii
Physcomitrium patens
Marchantia polymorpha
Anthoceros agrestis
Chara braunii
Chlamydomonas reinhardtii
Prasinoderma coloniale

Supp Fig 9

**A**

| | Y121 | ASG191-193 | V448 | S472 |
|---|---|---|---|---|
| *A. thaliana* PAL3 | — ELIR**Y**LNAG | — GTIT**ASG**DLVP | — KGAE**V**AMAS | — VESA**S**QHNQ |
| *A. suesica* PAL3A0A8T1Z666 | — ELIR**Y**LNAG | — GTIT**ASG**DLVP | — KGAE**V**AMAS | — VESA**S**QHNQ |
| *A. thaliana x arenosa* A0A8TCNU9 | — ELIR**Y**LNAG | — GTIT**ASG**DLVP | — KGAE**V**AMAS | — VESA**S**QHNQ |
| *A. lyrata* D7LXG7 | — ELIR**Y**LNAG | — GTIT**ASG**DLVP | — KGAE**V**AMAS | — VESA**S**QHNQ |
| *Arabis nemorensis* VVB13156.1 | — ELIR**Y**LNAG | — GTIT**ASG**DLVP | — KGGE**V**AMAS | — VESA**S**HHNQ |

**B**

| | *A. thaliana* PAL3 | *A. suesica* PAL3A0A8T1Z666 | *A. thaliana x arenosa* A0A8TCNU9 | *A. lyrata* D7LXG7 | *Arabis nemorensis* VVB13156.1 |
|---|---|---|---|---|---|
| *A. thaliana* PAL3 | | 90.8 | 99.8 | 94.4 | 87.7 |
| *A. suesica* A0A8T1Z666 | 95.4 | | 91.0 | 93.6 | 85.5 |
| *A. thaliana x arenosa* A0A8TCNU9 | 99.9 | 95.6 | | 94.5 | 87.9 |
| *A. lyrata* D7LXG7 | 97.3 | 97.3 | 97.4 | | 89.0 |
| *Arabis nemorensis* VVB13156.1 | 93.7 | 93.6 | 93.9 | 95.6 | |

Supp Fig 10

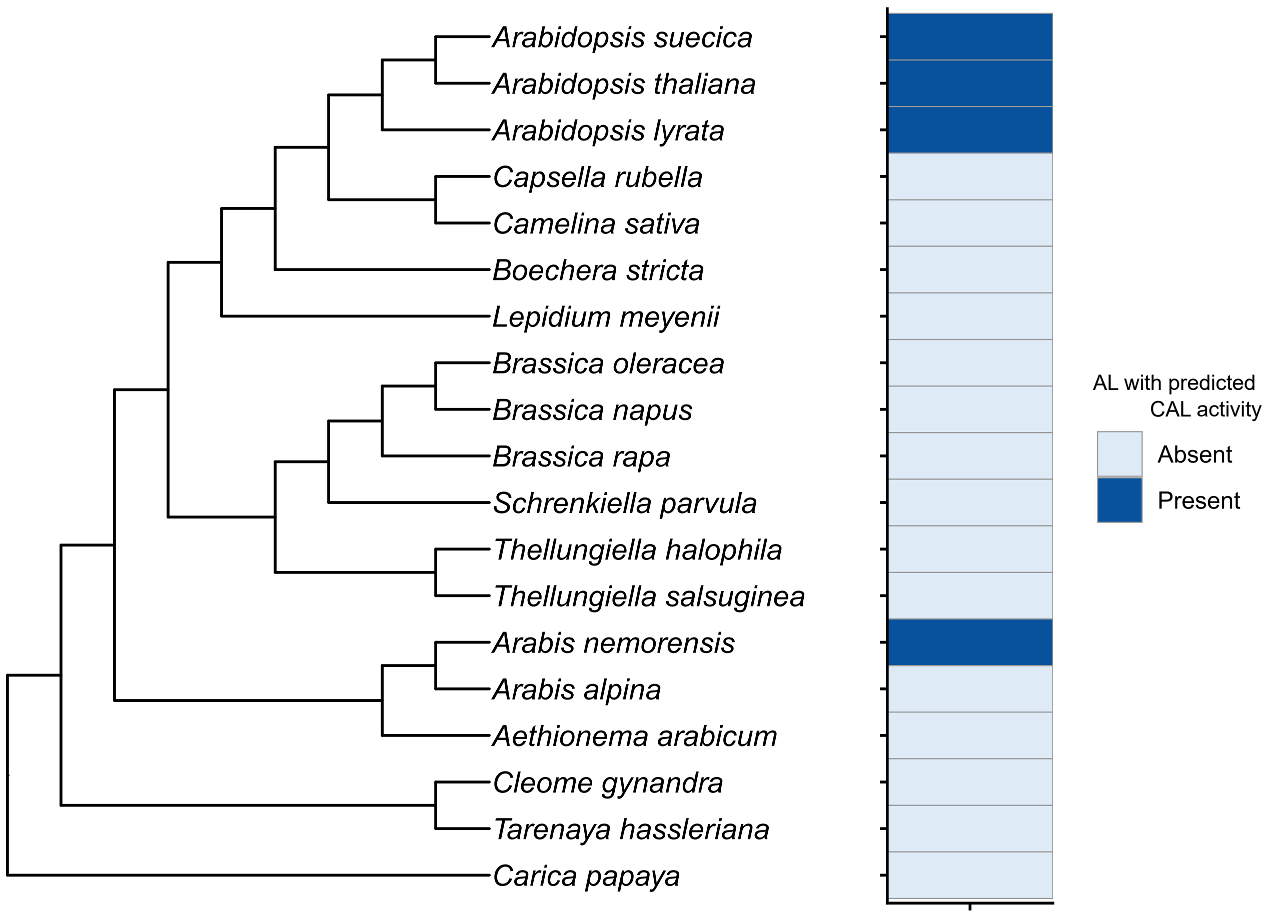
Arabidopsis suecica
Arabidopsis thaliana
Arabidopsis lyrata
Capsella rubella
Camelina sativa
Boechera stricta
Lepidium meyenii
Brassica oleracea
Brassica napus
Brassica rapa
Schrenkiella parvula
Thellungiella halophila
Thellungiella salsuginea
Arabis nemorensis
Arabis alpina
Aethionema arabicum
Cleome gynandra
Tarenaya hassleriana
Carica papaya
AL with predicted CAL activity
Absent
Present

Supp Fig 11

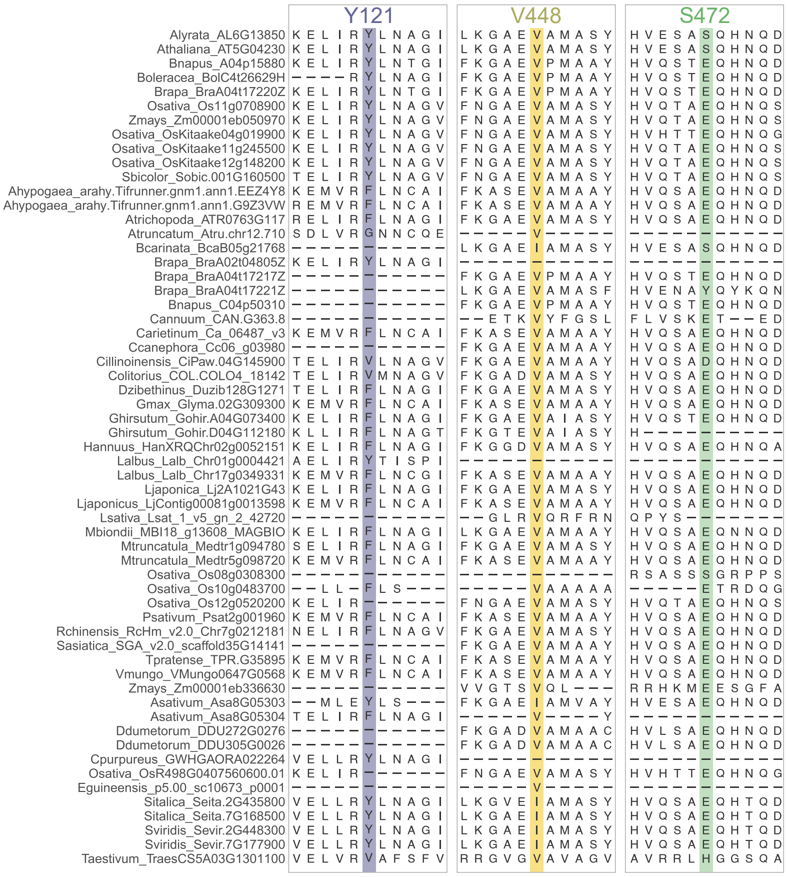


Supp Fig 12

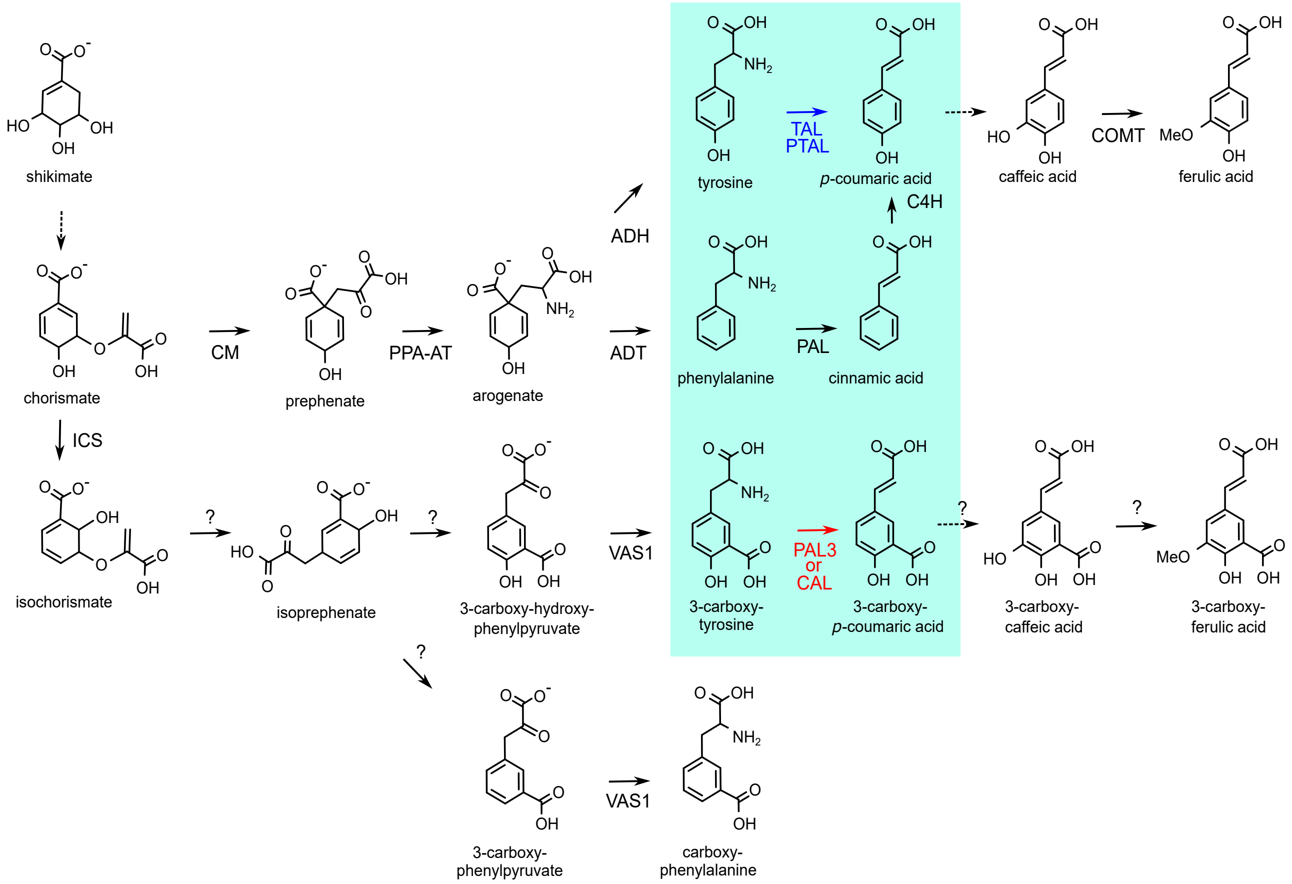
shikimate
chorismate
CM
prephenate
PPA-AT
arogenate
ADH
ADT
tyrosine
TAL
PTAL
p-coumaric acid
caffeic acid
COMT
ferulic acid
C4H
phenylalanine
PAL
cinnamic acid
ICS
isochorismate
?
isoprephenate
3-carboxy-hydroxy-phenylpyruvate
VAS1
3-carboxy-tyrosine
PAL3
or
CAL
3-carboxy-p-coumaric acid
3-carboxy-caffeic acid
3-carboxy-ferulic acid
3-carboxy-phenylpyruvate
carboxy-phenylalanine

Supp Fig 13

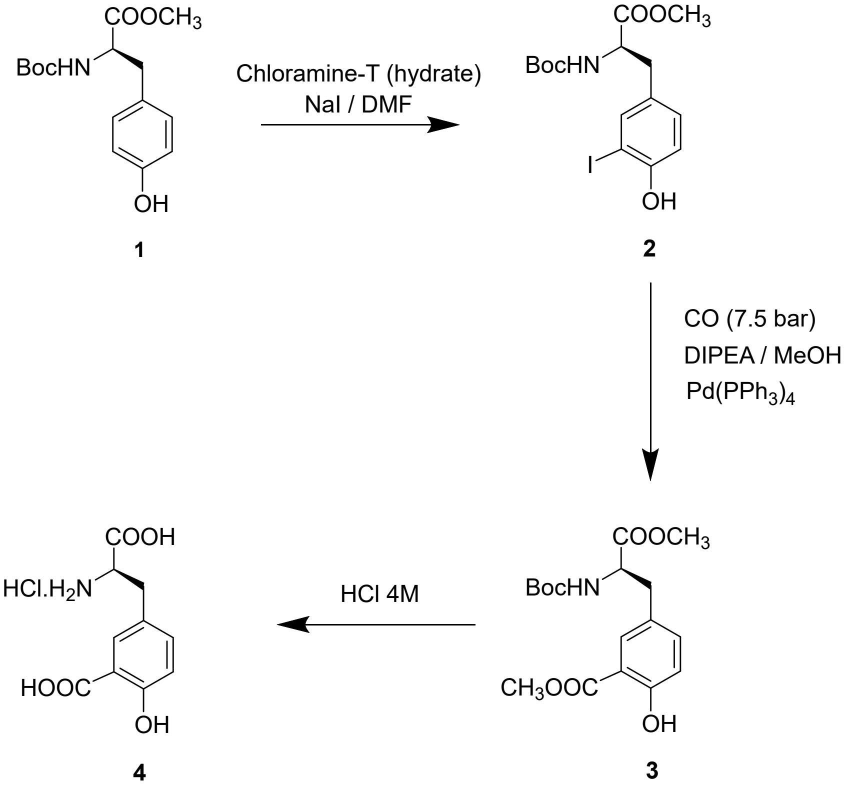
COOCH3
BocHN
OH
1
Chloramine-T (hydrate)
NaI / DMF
COOCH3
BocHN
I
OH
2
CO (7.5 bar)
DIPEA / MeOH
Pd(PPh3)4
COOCH3
BocHN
CH3OOC
OH
3
HCl 4M
COOH
HCl.H2N
HOOC
OH
4

Supp Fig 14
