## Supplementary material for "The Arabidopsis PAL3 is a carboxy-tyrosine ammonia lyase": Suppl Table 1

| Purpose | Nr | Description | Sequence |
|---|---|---|---|
| **Guide RNAs *PAL* mutagenesis** | 1 | gRNA PAL1 | 5'-GATCGCAGCCACTTGTCCAATGG-3' |
| | 2 | gRNA PAL2 | 5'-GTTGACTCTGACGAGCATGGCGG-3' |
| | 3 | gRNA PAL3 | 5'-GTTGCGGCAGTAGCGAGTGGAGG-3' |
| | 4 | gRNA PAL3 | 5'-GTTCCGTTATCCTATATCGCTGG-3' |
| | 5 | gRNA PAL4 | 5'-GTGATGGAGAGCATGAACCGAGG-3' |
| | 6 | gRNA PAL4 | 5'-GTGACGTGTCCCCGGCGCCGGGG-3' |
| **General primers** | 7 | FW primer Golden Gate Entry clone | 5'-GTGAGCGGATAACAATTTCACA-3' |
| | 8 | RV primer Golden Gate Entry clone | 5'-CGACGGCCAGGTAATACGACT-3' |
| | 9 | FW primer Golden Gate Expression clone | 5'-CGCACTCAGTCTTTCATCTACGG-3' |
| | 10 | RV primer Golden Gate Expression clone | 5'-GAGCTAGTTAGAATCTCAGGCCTTG-3' |
| | 11 | FW primer pGG vector | 5'-CAGGCTTTACACTTTATGCTTCCGGC-3' |
| | 12 | RV primer pGG vector | 5'-AGGGTTTTCCCAGTCACGACGTT-3' |
| | 13 | FW primer Gateway Recombination site attB1 | 5'-GGGGACAAGTTTGTACAAAAAAGCAGGCT-3' |
| | 14 | RV primer Gateway Recombination site attB2 | 5'-GGGGACCACTTTGTACAAGAAAGCTGGGT-3' |
| **Genotyping *pal* mutants** | 15 | FW primer to genotype pal1 crispr mutants | 5'-AGATTAACGGGGCACACAAG-3' |
| | 16 | RV primer to genotype pal1 crispr mutants | 5'-CGACACCGTTTTTGGTTCTC-3' |
| | 17 | FW primer to genotype pal1 crispr mutants | 5'-CTTAACCAACAATATTACAAATAAGCTCC-3' |
| | 18 | RV primer to genotype pal1 crispr mutants | 5'-TCCAACTAATAACAGCCACGTATTCTACC-3' |
| | 19 | FW primer to genotype pal2 crispr mutants | 5'-GAGATATACCGGGTCGGTGA-3' |
| | 20 | RV primer to genotype pal2 crispr mutants | 5'-GAAGTCCGGCGATGTAAGAG-3' |
| | 21 | FW primer to genotype pal3 crispr mutants | 5'-ACGGATCCCAAATTATATCAACCAGG-3' |
| | 22 | RV primer to genotype pal3 crispr mutants | 5'-CCACTCTTTCAGACATTCAAACAGAGG-3' |
| | 23 | FW primer to genotype pal3 crispr mutants | 5'-CGAAGATGACGACAGGTCAAACACG-3' |
| | 24 | RV primer to genotype pal3 crispr mutants | 5'-TCGACACTTCTTGATTCGGTTAGCCAC-3' |
| | 25 | FW primer to genotype pal4 crispr mutants | 5'-CCCTACTGGCTAGTAAAACAAAGC-3' |
| | 26 | FW primer to genotype pal4 crispr mutants | 5'-CTATTTAGCTGAATAAGTTGGTGTTTGG-3' |
| | 27 | RV primer to genotype pal4 crispr mutants | 5'-GCGATGTAAGAGAGAGGAACAAGG-3' |
| | 28 | FW primer to genotype pal4 crispr mutants | 5'-GAAGTTAGGAGGTGAGACTTTGACG-3' |
| | 29 | RV primer to genotype pal4 crispr mutants | 5'-TTGAGAAGCTTTGTAATTGCTTCGAG-3' |
| | 30 | FW primer to genotype pal4 crispr mutants | 5'-ACGGCCGAAGCTTTGAAAGGGAG-3' |
| | 31 | RV primer to genotype pal4 crispr mutants | 5'-GGTGATGGTGCCACGGAGAGGG-3' |
| **Cloning *PAL3* promotor** | 32 | FW primer outer promoter region PAL3 | 5'-GGATGAGTCCGATAGGGATTCCGATTAC-3' |
| | 33 | RV primer outer promoter region PAL3 | 5'-CATTCCAGTTAAGTGGGTCGCTCAATGC-3' |
| | 34 | FW primer inner promoter region PAL3 | 5'-GGGGACTGCTTTTTTGTACAAACTTGGATCACAAAATCTGACGGCAACC-3' |
| | 35 | RV primer inner promoter region PAL3 | 5'-GGGGACAACTTTGTATAGAAAAGTTGCCAACACACTGGTGAAGAAACCCAT-3' |
| **Cloning *PAL3*** | 36 | RV primer exon_1 PAL3 | 5'-CGTTCAAATACCTAATAAGCTCTTTTTGAAGAGC-3' |
| | 37 | FW primer exon_2 PAL3 | 5'-AAAGAGCTTATTAGGTATTTGAACGCCGGGATATTCG-3' |
| | 38 | RV primer exon_2 PAL3 | 5'-TCGCAGAGCGTAACCAAGAACATACCGATCTTGTTTAGG-3' |
| | 39 | FW primer exon_3 PAL3 | 5'-GATCGGTATGTTCTTGGTTACGCTCTGCGAACATCTCC-3' |
| | 40 | FW primer PAL3 CDS including attB1 | 5'-GGGGACAAGTTTGTACAAAAAAGCAGGCTTCATGGAGTTTCGTCAACCAAACG-3' |
| | 41 | RV primer PAL3 CDS including attB2 | 5'-GGGGACCACTTTGTACAAGAAAGCTGGGTGTTAGCAGATAGAAATCGGAGCACC-3' |
| **Cloning *PAL1*** | 42 | FW primer PAL1 CDS including attB1 | 5'-GGGGACAAGTTTGTACAAAAAAGCAGGCTTCATGGAGATTAACGGGGC-3' |
| | 43 | RV primer PAL1 CDS including attB2 | 5'-GGGGACCACTTTGTACAAGAAAGCTGGGTGTTAACATATTGGAATGGGAG-3' |
| **Mutagenesis of *PAL1*** | 44 | FW primer PAL1_F144Y | 5'-CAGAAGGAACTTATTAGATATCTTAACGC-3' |
| | 45 | RV primer PAL1_F144Y | 5'-CGAATATTCCGGCGTTAAGATATCTAATAAG-3' |
| | 46 | FW primer PAL1_I469V | 5'-GGAGCTGAGGTTGCAATGGCTT-3' |
| | 47 | RV primer PAL1_I469V | 5'-AAGCCATTGCAACCTCAGCTCC-3' |
| | 48 | FW primer PAL1_E493S | 5'-CATCTTGGTTATGTTGGCTTGCTGATTGAACATGG-3' |
| | 49 | FW primer PAL1_E493S | 5'-CTGTGACTAGCCATGTTCAATCAGCAAGCCAACATAACC-3' |
| **Mutagenesis of *PAL3*** | 50 | FW primer PAL3_Y121F | 5'-TCTTCAAAAAGAGCTTATTAGGTTCTTGAACG-3' |
| | 51 | RV primer PAL3_Y121F | 5'-CGAATATCCCGGCGTTCAAGAACCTAATAAGC-3' |
| | 52 | FW primer PAL3_Y121F | 5'-AAGAGCTTATTAGGTTCTTGAACGCCGGGATATTCG-3' |
| | 53 | RV primer PAL3_Y121F | 5'-CGTTCAAGAACCTAATAAGCTCTTTTTGAAGAGC-3' |
| | 54 | FW primer PAL3_V448I | 5'-AAGGCGCAGAAATCGCCATGGCTTCTTATTGCTCAGAGC-3' |
| | 55 | RV primer PAL3_V448I | 5'-AAGCCATGGCGATTTCTGCGCCTTTTAACCCGTAATC-3' |
| | 56 | FW primer PAL3_S472E | 5'-AACCATGTCGAAAGCGCTGAACAACACAATCAAGATG-3' |
| | 57 | RV primer PAL3_S472E | 5'-TTGTGTTGTTCAGCGCTTTCGACATGGTTCGTCACAGG-3' |
| **Cloning *ZmPAL1*** | 58 | FW primer exon1 outer ZmPAL1 | 5'-ATCATTGTTGTCGCCTCCTCCATTGCTCC-3' |
| | 59 | RV primer exon1 outer ZmPAL1 | 5'-CGTTGAGGTATCTGTAATATTTTCAGACC-3' |
| | 60 | FW primer exon2 outer ZmPAL1 | 5'-GGAGTACTTGATCACTCTGGGCTCTGG-3' |
| | 61 | RV primer exon2 outer ZmPAL1 | 5'-AACTGTGTGCCGTCAAGACACATGG-3' |
| | 62 | FW primer exon1 outer ZmPAL1 | 5'-TAGAAAGTAGCACACATGGGCACACATCAATCGG-3' |
| | 63 | FW primer exon1 inner ZmPAL1 | 5'-AAAAAGCAGGCTTCATGGAGTGCGAGACCG-3' |
| | 64 | FW primer ZmPAL1 CDS including attB1 | 5'-GGGGACAAGTTTGTACAAAAAAGCAGGCT-3' |
| | 65 | RV primer ZmPAL1 CDS including attB2 | 5'-GGGGACCACTTTGTACAAGAAAGCTGGGT-3' |
| **Cloning *OsPAL8*** | 66 | RV primer exon1 outer OsPAL8 | 5'-TTCATTCAGAACCACCCACCCAATGAACG-3' |
| | 67 | FW primer exon2 outer OsPAL8 | 5'-GTAATTAATTCGTTCATTGGGTGGGTGGTTCTG-3' |
| | 68 | RV primer exon2 outer OsPAL8 | 5'-TAAATAACGAGGGCATATGATTTCGTCGAG-3' |
| | 69 | FW primer exon2 inner OsPAL8 | 5'-AAAGAGCTGATCCGATACCTCAACGCCGGCGTCTTTGG-3' |
| | 70 | FW primer OsPAL8 CDS including attB1 | 5'-GGGGACAAGTTTGTACAAAAAAGCAGGCTTCATGGAGTGCGAGACCG-3' |
| | 71 | RV primer OsPAL8 CDS including attB2 | 5'-GGGGACCACTTTGTACAAGAAAGCTGGGTGTCAGATAATTGGCAGGGG-3' |
| **Cloning *BdPTAL*** | 72 | FW primer BdPTAL including attB1 | 5'-GGGGACAAGTTTGTACAAAAAAGCAGGCTTCATGGCGGGCAACGGAGCCATCTCG-3' |
| | 73 | RV primer BdPTAL including attB2 | 5'-GGGGACCACTTTGTACAAGAAAGCTGGGTGTTACTAGACGACGTTGATGGGCAGG-3' |
| **Cloning *AnPAL*** | 74 | FW primer AnPAL including attB1 | 5'-GGGGACAAGTTTGTACAAAAAAGCAGGCTTCATGGAATTCTGTCCCCCTAATAAGC-3' |
| | 75 | RV primer AnPAL including attB2 | 5'-GGGGACCACTTTGTACAAGAAAGCTGGGTGCTAACAGATTGGTATTGGGGCACCATTCC-3' |
